# Masserstein: robust linear deconvolution by optimal transport

**DOI:** 10.1101/2020.06.02.129858

**Authors:** Michał Ciach, Błażej Miasojedow, Grzegorz Skoraczyński, Szymon Majewski, Michał Startek, Dirk Valkenborg, Anna Gambin

## Abstract

A common theme in many applications of computational mass spectrometry is fitting a linear combination of reference spectra to an experimental one in order to estimate the quantities of different ions, potentially with overlapping isotopic envelopes. In this work, we study this procedure in an abstract setting, in order to develop new approaches applicable to a diverse range of experiments. We introduce an application of a new spectral dissimilarity measure, known in other fields as the Wasserstein or the Earth Mover’s distance, in order to overcome the sensitivity of ordinary linear regression to measurement inaccuracies. Usinga a data set of 200 mass spectra, we demonstrate that our approach is capable of accurate estimation of ion proportions without extensive pre-processing required for state-of-the-art methods. The conclusions are further substantiated using data sets simulated in a way that mimics most of the measurement inaccuracies occurring in real experiments. We have implemented our methods in a Python 3 package, freely available at https://github.com/mciach/masserstein.

## 1 Introduction

A frequently encountered task in mass spectrometry is the quantification of signal corresponding to a particular set of ions. From an algorithmic point of view, the most challenging parts of this problem are the separation of the signal from the noise and the separation of overlapping isotopic envelopes. Numerous approaches have been developed in order to tackle this problem in the context of specific experiments. These include the estimation of reaction rates in ETD fragmentation [1]; quantification of polymer chain lengths and compositions [2, 3]; annotation of MS^2^ spectra in data independent acquisition label-free quantification experiments [4]; studies of fragmentation of aliphatic diselenides and selenosulfenates [5, 6], and studies of protein deamidation and 18 O labelling [7].

Despite the abundance of different algorithmic approaches and software tools, a common theme is the approximation of an experimentally observed spectrum by a set of reference spectra. Therefore, from a mathematical point of view, all the described problems can be expressed with a single equation:

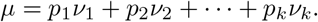

Here, *μ* is the observed spectrum, *v*_*i*_’s are the reference spectra of the ions in question, and *p*_*i*_’s are the unknown proportions of the latter. Since, in mass spectrometry, the reference spectra are most often predicted using computational methods, we refer to them as *theoretical* spectra. A common theme of different approaches to linear deconvolution is the assumption that the reference spectra are known. Therefore, from the methodological point of view, this problem is an exact opposite of molecule identification, where the task is to identify the identity of ions, but not their quantities.

The equation presented above has appeared e.g. in [4], where the authors have used it to annotate data independent acquisition label-free quantification experiments. Since the experimental spectrum is expressed as a linear combination of the theoretical ones, this problem has been termed *linear deconvolution* [4]. An example of a linear deconvolution is shown in Fig. 1, which illustrates a linear deconvolution of a mass spectrum consisting of overlapping isotopic envelopes of human hemoglobin subunits *α* and *δ*.

**Figure 1:**
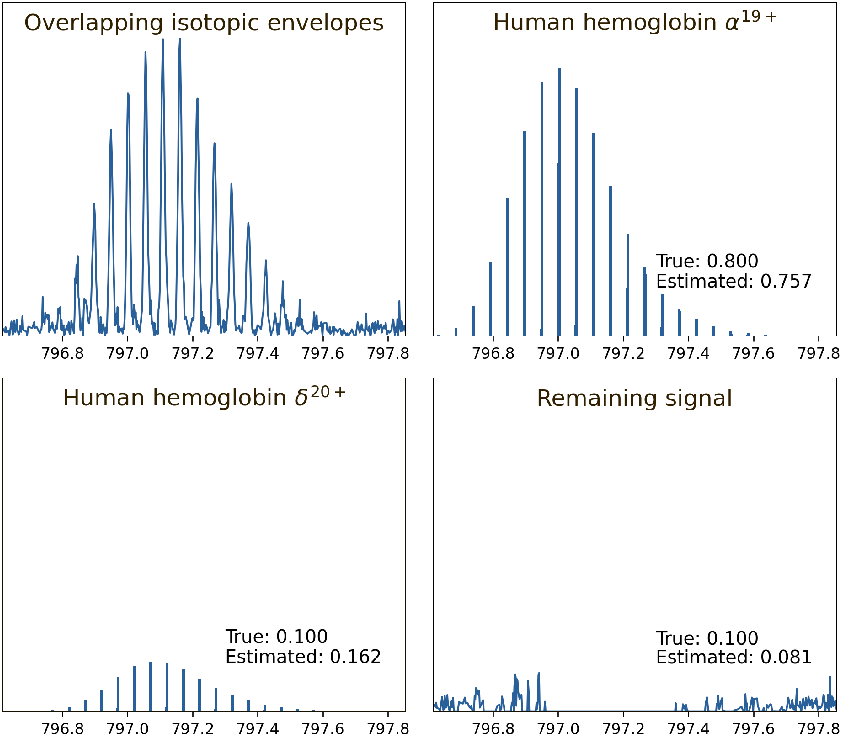
An example of deconvolution of a fragment of a simulated human hemoglobin ESI MS^1^ spectrum. The fragment is deconvolved into signals of *α* 19+ and *δ* 20+ subunits and the remaining background noise using the method presented in this article. The proportions were estimated directly from the top-left spectrum, without any additional preprocessing such as peak picking or denoising.

Unfortunately, the currently used terminology is somewhat misleading. The problem of estimating abundances of convolved isotopic envelopes is known in the literature under several names, most common ones being the *resolving of isobaric interferences* and *deconvolution*. On the other hand, the latter term itself has several meanings which have little in common. One of the most common ones is converting m/z values into masses, as exemplified by the popular MaxEnt algorithm [8], also referred to as the *charge deconvolution*. One of the major differences between the charge and the linear deconvolution is that the former usually does not assume the knowledge of the reference spectra. In this article, we use the term deconvolution solely to refer to the estimation of an ion’s signal in the presence of overlapping isotopic envelopes and/or background noise.

In the case when different isotopic envelopes do not overlap, the background noise is small, and there is only a handful of molecules of interest, linear deconvolution can easily be performed manually by integrating selected regions of the experimental spectrum. However, algorithmic approaches need to be employed when there is a considerable overlap of isotopic envelopes, the signal-to-noise ratio is small, or thousands of molecules need to be analyzed in a high-throughput setting.

A common approach, found e.g. in the specter [4] and the masstodon [9] tools, is to perform a linear regression where the experimental spectrum is treated as the dependent variable and the theoretical ones as independent variables. Mathematically, this technique minimizes the Euclidean distance between the experimental spectrum and a linear combination of the theoretical ones. A similar technique, the L1 regression (also known as the least absolute deviation regression), is sometimes used [3]. Another approach is a sequential subtraction of estimated signal from the experimental spectrum, as exemplified by the THRASH algorithm [10].

The performance of such methods is hindered by a series of problems. Most of them arise from the fact that they are based on pointwise comparison of spectra, that is, they compare peaks with the same m/z values. However, unlike the theoretically predicted spectra, experimental ones have limited resolution and accuracy. Because of this, peak picking of the experimental spectrum is required, which is often imperfect (such as the conventional peak centroiding) or computationally expensive (such as the continuous wavelet transform approach [11]). The experimentally obtained peaks virtually never match the theoretical ones perfectly due to accuracy limits of the instrument and numerical errors of peak-picking procedures.

### Our contribution

The goal of this work is to further investigate the problem of linear deconvolution of spectra. We develop an algorithm that allows for a simultaneous deconvolution and denoising of spectra while being free from the limitations of methods based on linear regression. In particular, the method is robust against measurement inaccuracies and numerical errors of peak picking algorithms, while being resilient to interfering signals coming from chemical impurities and background noise.

The main novelty in our approach to linear deconvolution is the use of the Wasserstein distance, which is a recently proposed approach to processing mass spectra [12]. This distance, based on the theory of the optimal transport, is naturally robust to uncertainties in m/z measurements and different resolutions of the compared spectra. In particular, to our knowledge, this is the first algorithm that is capable of explaining an experimental spectrum in profile mode using a set of computationally generated, infinitely resolved theoretical spectra. On the other hand, when the experimental spectrum is analyzed in centroid mode, our algorithm does not require peak matching, and is therefore capable of utilizing the full information in both the experimental and the theoretical spectra.

We tackle the problem of linear deconvolution in its abstract form instead of focusing on a particular type of experiment. This way, while being mathematically sound, our methods retain their full scope of applicability, from analytical chemistry to metabolomics to proteomics to synthetic polymer science. They are not limited to any single type of mass spectrometer or pre-processing software. The mathematical foundations stay the same regardless whether an FTICR, TOF, or quadrupole instrument is used.

Accordingly, instead of presenting an application of our linear deconvolution algorithm to any particular experiment, we test its performance on a custom-made data set consisting of 200 repeated measurements of the same set of compounds. This allows us to assess both the accuracy and variance of the estimation of ion proportions. We further confirm our results using simulated data sets, where we take into account several measurement inaccuracies occurring naturally in mass spectra.

The deconvolution algorithm presented in this work has been implemented in the Python 3 language, and is freely available as a set of command-line applications and a Python 3 package `masserstein` at https://github.com/mciach/masserstein.

### Structure of the article

First, we give an introduction on the Wasserstein distance, an optimal transport-based approach to comparison of spectra. Next, we describe the linear deconvolution problem in detail and derive a deconvolution algorithm based on the Wasserstein distance, which is the main contribution of this work. Finally, we assess the performance of the algorithm on a set of 200 experimental spectra as well as on computationally generated datasets.

## 2 Methods

As described in the introduction, naturally occurring measurement inaccuracies hinder the performance of the common approaches to the estimation of relative abundances of ions in mass spectra. In order to overcome these problems, a spectrum dissimilarity measure based on the theory of optimal transport was recently investigated, with the aim to develop a linear deconvolution method that allows to compare spectra with different resolutions and is robust to measurement errors in the mass domain [12]. The measure is known in the field of probability theory as the Wasserstein metric [13], and in the computer science community as the Earth Mover’s distance [14].

The idea behind the measure is to transport the ion current from one spectrum onto the other one, and quantify the distance that the current needs to travel. The dissimilarity between the two spectra is equal to the minimal distance in m/z domain that needs to be traveled in order to fully transform one spectrum into the other. In particular, the Wasserstein distance between MS^1^ spectra of two ions with the same charge is approximately equal to the molecules’ mass difference. This interpretation allows to consider as fairly similar those spectra for which the Wasserstein distance is less than one hydrogen mass. Examples of the values of this distance are shown in Fig. 2. In the right panel, we consider a pair of corresponding spectra in profile and centroid mode. Even though such spectra are incomparable using conventional measures such as the Euclidean distance, the Wasserstein distance between them has a very small value of 0.032 Da. This value reflects the different resolutions of those spectra. The left panel presents a pair of spectra with no matching peaks. Pointwise measures such as correlation do not capture any similarity between them (note that in some applications this may be a desirable phenomenon). The Wasserstein distance, on the other hand, again has a small value of 0.078 Da, indicating that those spectra are very similar in terms of the m/z differences of peak positions. The fact that the peaks do not match, however, is still reflected by the Wasserstein distance, since it is over twice as large as between the spectra in the right panel. The exact method of calculation of this distance and some of its properties when applied to mass spectra are discussed further in the article.

**Figure 2:**
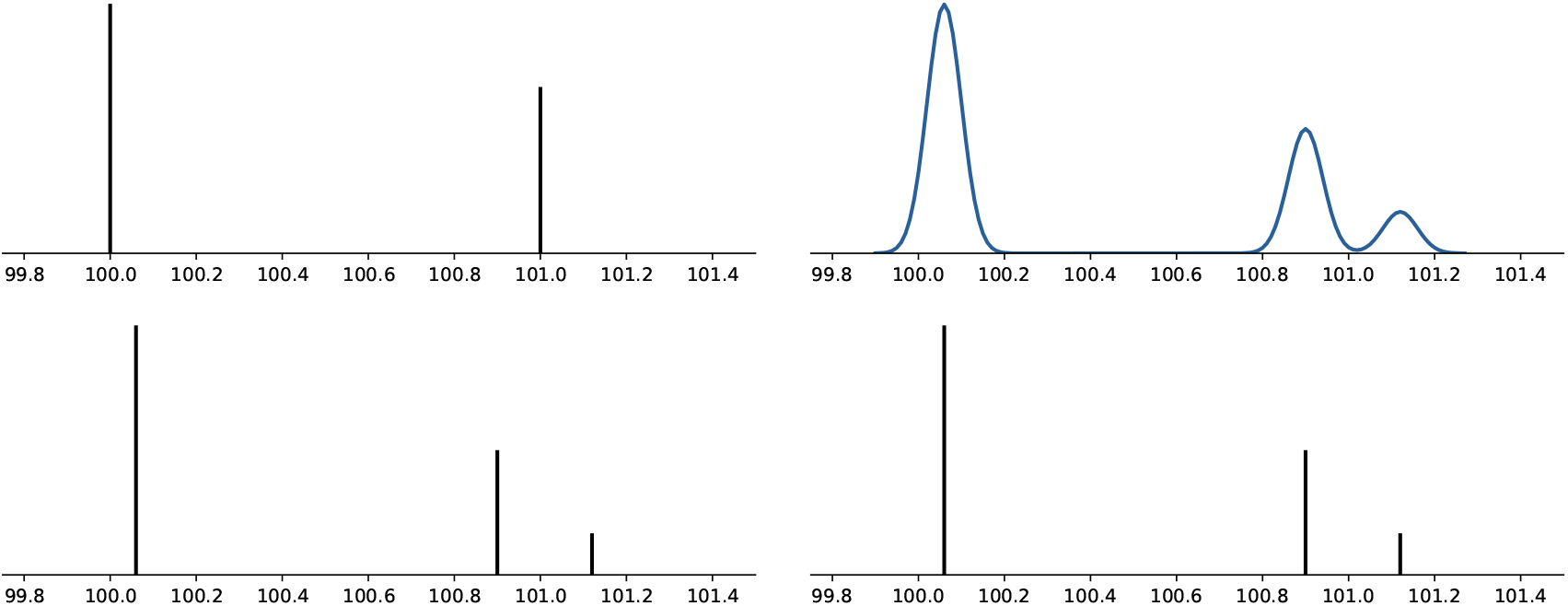
Example values of the Wasserstein distance between mass spectra. Left: Two spectra in centroid mode, with the Wasserstein distance between them being equal to 0.078 Da. Right: A spectrum in profile mode and a corresponding centroided one, with the Wasserstein distance equal to 0.032 Da. Both values, being less than 1 hydrogen mass, indicate a fairly high degree of similarity, even though no peaks match in the first example and the spectra are in different modes (profile vs centroid) in the second one.

An alternative interpretation of the distance is the minimal amount of distortion such as shifting, broadening and narrowing of peaks that is required to transform one of the spectra into the other. This approach is naturally robust to small distortions in the m/z or intensity measurements, and has no requirements as to the accuracy or resolution of the measurement. In particular, to the best of our knowledge, it is the only similarity measure capable of an accurate comparison of profile and centroided spectra.

In the following subsections, we give a short introduction to the mathematical formalism behind the Wasserstein distance and show several examples that illustrate its properties related to mass spectra. Next, we develop a deconvolution algorithm which is the main contribution of this work. The algorithm improves upon the original one published in [12] by allowing to discard signal from the experimental spectrum during deconvolution, thus allowing for simultaneous quantification and denoising. This makes the method robust to background noise, and makes it applicable to a broader class of experimental spectra. We also improve on the computational complexity of the original algorithm.

### 2.1 The Wasserstein distance

Let *μ* and *v* be any two mass spectra to be compared. We aim to transport all the signal from one spectrum to the other and quantify the minimal amount of distance in the mass domain over which the signal needs to be transported. We do not differentiate between a source and a target spectrum, so that the final distance is symmetrical—in other words, we will assume that transforming *μ* into *v* inflicts the same cost as transforming *v* into *μ*.

We treat mass spectra as functions defined on the real line ℝ, so that *μ*(*x*) is the intensity at the m/z value *x* in the spectrum *μ*. We assume that all the considered spectra are normalized by their total ion current, so that the peak intensities of each spectrum sum up to 1.

Let *γ*(*x, y*) be the amount of signal transported between the point *x* in spectrum *μ* and the point *y* in spectrum *v*. The function *γ* is referred to as a *transport plan*. Any transport plan needs to satisfy the following properties:

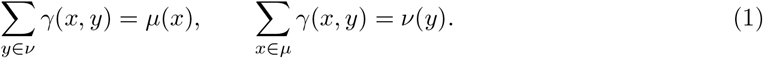

The first of those properties means that all the signal intensity *μ*(*x*) needs to be transported some-where into the spectrum *v*. Similarly, the second property means that the intensity *v*(*y*) is fully filled by the ion current coming from the spectrum *μ*. Naturally, the transport plan also needs to be non-negative, *γ*(*x, y*) ≥ 0, as we cannot transport negative intensity. In turn, any function satisfying the above properties is a valid transport plan. We denote by Γ the space of all possible transport plans.

The cost of a given transport plan is the total distance traveled by the ion current. This is calculated by multiplying the distance between the points *x* and *y* by the amount of signal traveling between them, and summing over all peaks in both spectra:

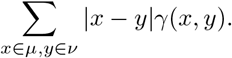

Note that the dependence of this sum on the spectra *μ* and *v* is only implicitly expressed through the function *γ* and equations (1).

We assume that the transport is not directed, so that transporting the intensity in the direction of increasing mass inflicts the same cost as in the other direction. This is formally expressed by the absolute difference between the points in the above equation.

The Wasserstein metric [13, 15] between the two spectra, *W* (*μ, v*), is defined as the minimal cost of transport over all possible transport plans from the space Γ:

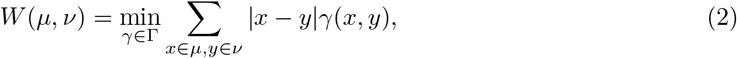

The function *W* defined this way satisfies the mathematical properties of a distance function [13], that is, non-negativity *W* (*μ, v*) ≥ 0, symmetry *W* (*μ, v*) = *W* (*v, μ*) and the triangle inequality *W* (*μ, v*) ≤ *W* (*μ, ζ*) + *W* (*ζ, v*).

Since the distance is symmetric, there is no designated *source* spectrum of the transported signal, nor a *target* spectrum to which the signal is transported. However, when depicting transport plans for *W* (*μ, v*), we adopt a convention that the signal is transported from *v* to *μ*.

Although the formulation of the distance may seem baffling, it turns out that under reasonable assumptions the algorithm to compute *W* (*μ, v*) has a linear time complexity [15]. The only requirement for this is that the input spectra are available as peak lists sorted by their m/z value, which is the case in most situations. Therefore, from the computational point of view, the Wasserstein metric is equally efficient as the Jaccard score or the Euclidean distance.

The algorithm to compute the Wasserstein distance between a pair of spectra is based on the following theorem [16, 15]:

#### Theorem 1

*Let μ and v be two probability measures on the real line* ℝ. *Let M and N be the cumulative distribution functions of μ and v respectively. Then,*

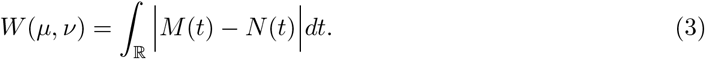

The above theorem can be applied to spectra normalized by their total ion current, since such spectra can be interpreted as probability measures. It captures both the case of centroided and profile spectra. In the case of the former, however, the cumulative distribution functions are step functions, and the formula can be considerably simplified.

#### Theorem 2

*Let μ, v be two centroided mass spectra normalized by the total ion current. Let S* = {*s*_1_, *s*_2_, … , *s*_*n*_} *be the list of all distinct ma*L*sses in both spectra. Let M and N be the cumulative* distribution functions of *μ* and *v*, i.e. *M*(t) = Σ_*x*≤*t*_ *μ*(*x*). *Then*,

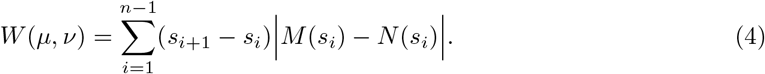

Formula (4) admits a simple interpretation. Observe that *M* (*s*_*i*_) − *N* (*s*_*i*_) is the difference in ion current on the left hand side of point *s*_*i*_, which needs to be transferred either to or from the point *s*_*i*+1_ in order to achieve balance.

In order to illustrate the computation of the Wasserstein distance and make the concept more intuitive to the reader, we present two worked examples below.

#### Example 1.

Consider two abstract spectra, *μ* and *v*, such that *μ* is concentrated at 100 Da (i.e. *μ*(100.5) = 1) and *v* is distributed evenly over 98, 99, 100, 101 and 102 Da (i.e. *v*(98) = *v*(99) = *v*(100) = *v*(101) = *v*(102) = 0.2). Obviously, those spectra do not correspond to any actual ions. They only serve for an easy illustration of the computation of the optimal signal transport. The procedure is illustrated in Fig. 3.

**Figure 3:**
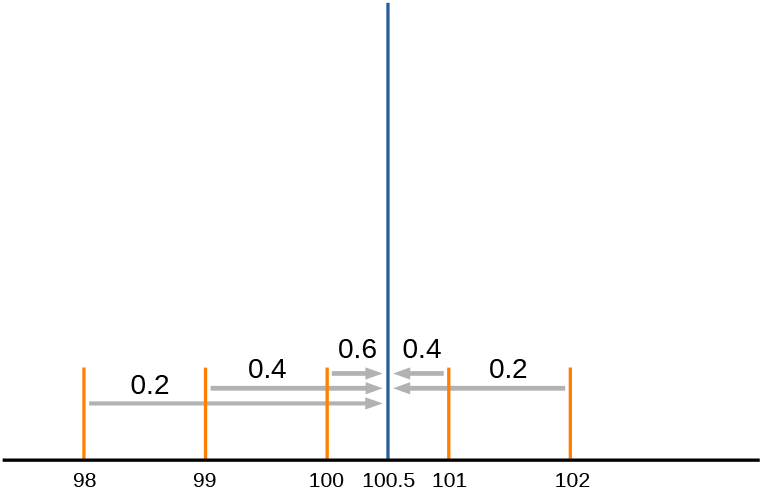
An example of an optimal transport scheme between two abstract spectra *μ* and *v*, depicted on a single graph in blue and orange respectively. The signal of the spectrum *μ* is concentrated at the point 100.5, while *v* is distributed evenly at five points from 98 to 102. Grey arrows depict the transport of the ion current. Numbers above arrows show the proportions of the ion current flowing between neighboring peaks.

The cumulative distribution function of *μ* and *v* are given by

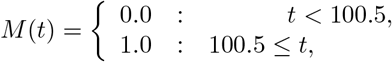

and

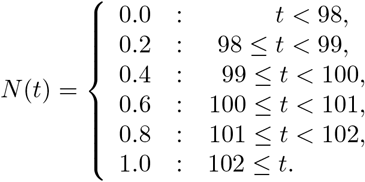

The list of all distinct masses, *S*, is now equal to (98, 99, 100, 100.5, 101, 102), and the difference between the cumulative distribution functions, *N*(*t*) − *M*(*t*), indicating the ion current imbalance at point *t*, is equal to

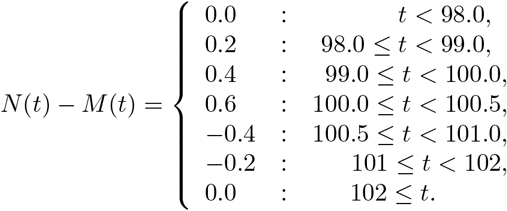

From the above equation, we can read out that 0.2 of the signal is transported from the point 98.0 to 99.0; 0.4 of the signal is transported from 99.0 to 100.0; 0.6 from 100.0 to 100.5. Next, as the sign of the imbalance changes, so does the direction of transport between neighboring points, and so 0.4 of the signal is transported from 101.0 to 100.5, and 0.2 of the signal from 102 to 101. Finally, the ion currents of both spectra balance out at the point 102.

The final distance can be computed by taking the absolute values of the ion current imbalance and multiplying them by the distance travelled, so that *W* (*μ, v*) = 0.2 1+0.4 1+0.6 0.5+0.4 0.5+0.2 1 = 1.3 Da.

#### Example 2.

Consider a spectrum *μ* concentrated at 100 Da, and a profile spectrum *v* consisting of a single Gaussian peak centered at 100 Da with a standard deviation *σ*. We therefore have *μ*(100) = 1 and

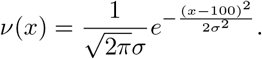

Let Φ be the cumulative distribution function of the standard Gaussian random variable:

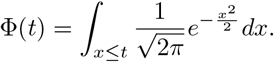

The CDF of *v* is then given by *N*(*t*) = Φ((*t* − 100)/*σ*). The exact value of the Wasserstein distance between *μ* and *v* is then equal to

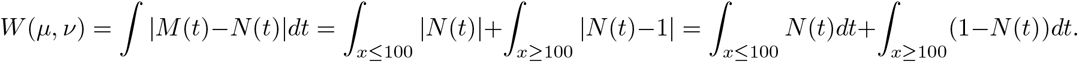

After plugging in Φ in the above integrals and substituting the variables, one arrives at

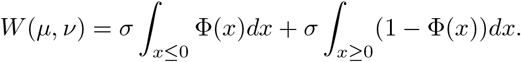

Using the relation Φ(−*x*) = 1 − Φ(*x*), we get

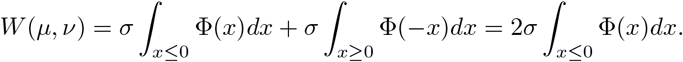

As the antiderivative of Φ(*x*) is *x*Φ(*x*) + *ϕ*(*x*) + *C*, where *ϕ* is the density of a standard Gaussian variable, we arrive at

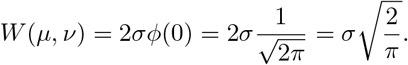

Therefore, we obtain a simple formula for the distance between a centroided spectrum and a corresponding profile spectrum with Gaussian peaks of standard deviation *σ*. This result explains the small distance between spectra in the right panel of Fig. 2.

### Handling profile spectra in practice

Although, in principle, profile spectra are continuous functions, they are usually represented as finite lists of mass and intensity pairs. It turns out that such lists can be simply treated as centroid spectra when the Wasserstein distance is computed. In that case, Theorem 2 gives an accurate approximation of the cost of the optimal transport plan.

Assume we are given a finite list of mass and intensity pairs, (*s*_*i*_, *I*_*i*_) for *i* = 1, 2, … , *n*, approximating a profile spectrum *μ*. Assume as well that we have a constant spacing between consecutive intensity measurements, so that *s*_*i*+1_ − *s*_*i*_ = 1/*n* for *i* = 1, 2, … , *n* − 1. Treating *μ* in the same way as a centroided spectrum, we compute its cumulative distribution function as 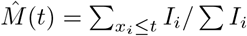. The true cumulative distribution function of *μ*, on the other hand, is given by *M*(*t*) = ∫_*x*≤*t*_ *μ*(*x*)*dx*/ ∫ *μ*(*x*)*dx*. Now, we have

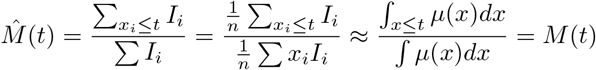

The assumption of an uniform signal sampling (i.e. a constant spacing between consecutive measurements) is crucial for the approximation to work. When this is not satisfied, a spectrum needs to be resampled prior to computing the distance. For a practical example of this procedure, see Section 3. The resampling algorithm used in this study, based on piecewise-linear interpolation of profile spectra, is presented in section 4.4 of the Appendix.

### 2.2 Linear deconvolution

In this section, we formally define the problem of linear deconvolution [4]. We consider an observed spectrum, denoted *μ*, and a library of expected spectra, denoted *v*_1_, … , *v*_*k*_. The expected spectra may come from theoretical simulations as well as experimental measurements, or a combination of both.

As before, we assume that all spectra are normalized by their total ion current. Let *p* = (*p*_1_, *p*_2_, … , *p*_*k*_) be a vector of non-negative *weights* or *proportions* such that *p*_1_ + … + *p_k_* ≤ 1. Define *v*_*p*_ to be a linear combination of the expected spectra with weights *p*, i.e.

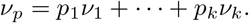

We will call *v*_*p*_ a *theoretical model* or a *model spectrum*, in analogy to a model matrix used in an ordinary least squares linear regression. Note that the *v_p_*’s intensity sums up to *p*_1_ + … + *p*_*k*_, which may be less than 1. The inequality is allowed to account for the fact that *v*_*p*_ may not explain all of *μ*’s signal.

We assume that the observed spectrum can be approximated by the model spectrum with some additional chemical and/or background noise *ε*, so that for any m/z value *x* we have

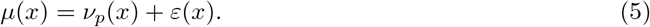

Note that because of the assumed normalization of the spectra, the total signal of *v*_*p*_ is the proportion of *μ* that is explained by the model.

The problem of linear deconvolution is defined as finding proportions which minimize some chosen distance function *d* between the observed spectrum and the combination of the expected spectra:

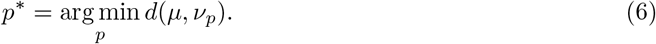

For example, in the commonly used least-squares approach, *d*(*μ*, *v*_*p*_) = Σ_*x*∈*S*_(*μ*(*x*) − *v*_*p*_(*x*))^2^, where *S* is the set of observed peak masses. As described in the introduction, this particular distance is sensitive to different resolutions and mass measurement errors, which makes it necessary to centroid the spectra and set a mass window for peak matching. The Wasserstein distance, on the other hand, is naturally robust to small errors in the mass measurement. Therefore, it is a natural candidate for a distance measure to be used in a linear deconvolution algorithm. By coupling deconvolution with a denoising procedure, we make it robust to errors in the intensity domain as well.

### Deconvolution by optimal transport

If all the signal of *μ* can be explained by the model spectrum *v*_*p*_, the deconvolution problem can be expressed as finding proportions *p*\* such that

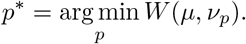

This optimization problem has been shown to yield accurate results when the only differences between *μ* and *v*_*p*_ are caused by mass measurement errors and differing resolution [12]. However, experimentally obtained spectra most often contain signals which are not theoretically predicted, like chemical or background noise. Such signals strongly disturb the optimal transport plan, leading to incorrect estimation of the proportions *p*\*. This is because the Wasserstein distance requires both spectra to have equal amounts of intensity, enforcing *p*_1_ + … + *p*_*k*_ to be equal to 1.

To account for the additional signal in *μ*, we extend the m/z axis by adding an auxilliary point *ω* onto which such signal can be transported. We assume that the cost of transporting signal from *μ* to *ω* does not depend on the m/z value of the signal (equivalently, that *ω* is equidistant to all points on ℝ). The idea behind this approach is visualized in Fig. 4. We denote as *κ* the cost of transporting signal from *μ* to *ω*(equivalently, the distance between *ω* and any point on ℝ). Since *κ* is interpreted as the cost of removing noise from *μ*, we refer to it as the *denoising penalty*.

**Figure 4:**
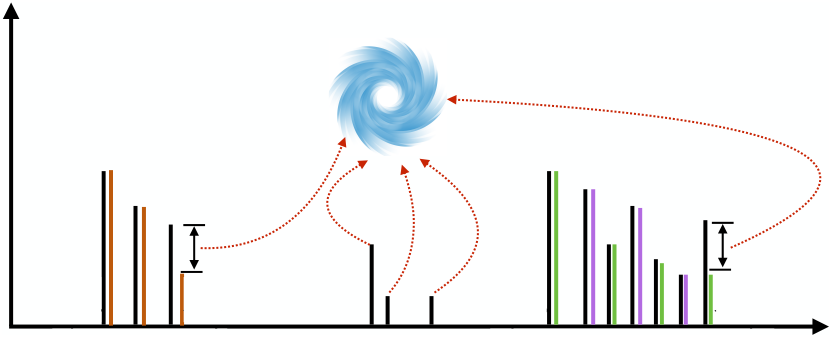
An illustration of estimation of proportions coupled with denoising. An experimental spectrum (black) is explained by a set of three theoretical spectra (orange, green and magenta). Excess signal from the theoretical spectrum, occurring due to background noise or sample impurities, is transported onto an auxilliary point *ω*, represented as the vortex.

The combined deconvolution-denoising problem is defined as

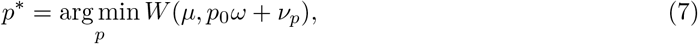

where *p*_0_ = 1 − *p*_1_ – … – *p*_*k*_ is the amount of the unexplained signal in *μ* transported onto *ω*.

Note that the denoising penalty *κ* admits a physical interpretation in terms of m/z units. Transporting intensity from the experimental to the model spectrum over a distance larger than *κ* is more costly than removing it by transporting it to *ω*. The penalty can therefore be treated as a maximum distance over which the transport is feasible. This allows for some intuition behind the optimal values of this parameter: the maximum feasible transport distance should be set as the smallest value that allows to match corresponding theoretical and experimental peaks. The instrument accuracy (in Dalton units) is therefore an example of a reasonable value for *κ*. In practice, however, the choice of this parameter is more complicated. This is because, apart from the instrument accuracy, there are several factors that influence the distance between experimentally observed peaks and their theoretical counterparts, including, but not limited to, the resolving power. Therefore, usually the results for several different values of *κ* need to be inspected manually. This issue is discussed in more detail in the subsequent sections.

A major advantage of our approach, as opposed to matching peaks by mass windows in methods based on linear regression, is that *κ* does not set a hard threshold on the transport distance, allowing for more flexibility in the choice in this parameter. In some cases it may be beneficial to transport the signal over distances larger than *κ*, while in other cases interfering signal is removed regardless of its proximity to theoretical peaks. Specifically, whether a given signal is removed depends not only on it’s distance from the nearest theoretical peaks, but also on the shapes of the theoretical isotopic envelopes—in contrast to methods based on linear regression, in which a signal is always incorporated when it’s sufficiently close to a theoretical peak. This phenomenon is illustrated by the following example.

#### Example 3.

Consider a theoretical spectrum *v* consisting of *n* peaks, and let the *i*-th peak be at m/z *m*_1_ +*i/q* for some *m*_*i*_ ≥ 0, and let it have intensity *a*_*i*_, where *a*_1_ + *a*_2_ + … + *a*_*n*_ = 1. This models a low-resolution theoretical spectrum of an ion with charge *q* and with the monoisotopic peak at *m*_1_. Since the proportions *a*_*i*_ are arbitrary, this model encompasses all possible single-ion low-resolution spectra, allowing us to obtain a general result.

Let the experimental spectrum *μ* be equal to the theoretical one scaled by 1 − *ϵ*, with an additional noise peak in location *x* ≤ *m*_1_ with intensity *ϵ* (see Fig. 5). We will investigate which values of *κ* allow to correctly identify the additional peak as noise, and return the proportion of *v* as 1 − *ϵ* instead of 1. To this end, we will compare the Wasserstein distance *W*(*μ, v*) with the cost of denoising *κ*. Note that, since the spectra are identical save for the noise peak, *κ* is the full cost of deconvolution when this peak is removed.

**Figure 5:**
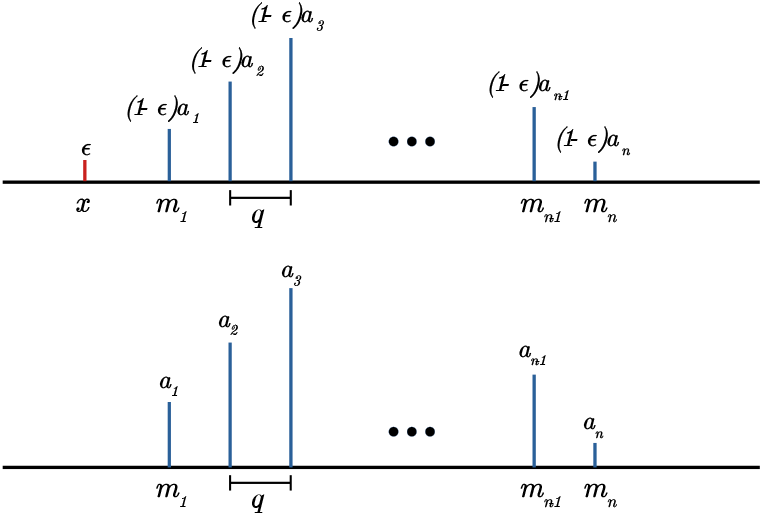
An example pair of experimental and theoretical spectrum, where the former (top) is equal to the latter (bottom) plus an additional noise peak.

Let *A*_*i*_ be the cumulative intensity of the theoretical spectrum *v*. The Wasserstein distance *W*(*μ, v*) is equal to

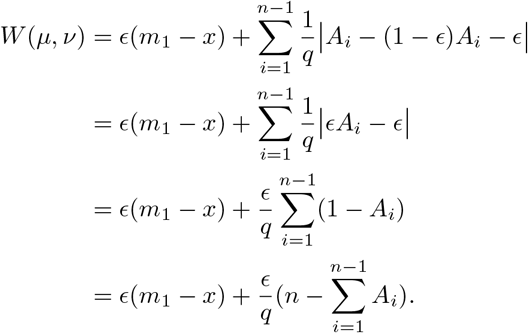

Now, note that 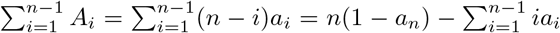, where the first equality comes from counting the number that each of the *a*_*i*_ variables occur in the sum, and the second one follows from the fact that the intensities sum to 1. This can be further simplified by noting that 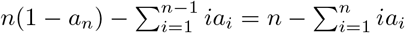, where we included the term *na*_*n*_ in the sum.

Now, note that since the intensities are normalized,the average mass of the spectrum *v*, denoted as 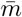, is equal to 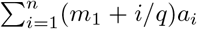. This allows us to express 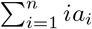 simply as 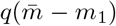, i.e. the distance between the average and the monoisotopic mass multiplied by the charge. Plugging this into the equation for *W*(*μ, nu*), we get a simple formula

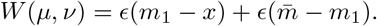

The cost of removing the noise peak will be lower than the cost of such transport whenever 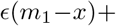 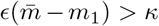. It follows that the influence of the noise peak on the estimated proportions depends not only on it’s proximity to the signal, *m*_1_ − *x*, as is the case in methods based on linear regression, but also on the width of the isotopic envelope, 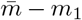. Note that the latter term is always positive for spectra with more than one peak. Therefore, the shape of the isotopic envelope facilitates the detection of noise peaks by the linear deconvolution procedure based on the Wasserstein distance.

### Computation of the optimal proportions

The formulation of the deconvolution-denoising problem (7) is well suited for theoretical analysis. However, it does not show how to obtain the optimal proportions in practice. Below, we show an equivalent formulation that is better suited for implementation and practical applications. The proof that the two formulations are equivalent, based on Theorem 2, has been relegated to the Appendix.

Let *M*(*t*), *N*_*j*_(*t*) be the cumulative distribution functions of *μ* and *v*_*j*_ respectively. The model spectrum 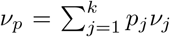 is a spectrum a with cumulative distribution function equal 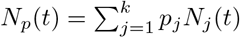.

Let *g*(*s*_*i*_) for *i* = 1, 2, … , *n* be the amount of *μ*’s signal at the point *s*_*i*_ that is labeled as noise, i.e. transported onto *ω*. Note the distinction between *g* and *ω*: the latter is an auxilliary spectrum, therefore it’s a concept analogous to *μ* and *v*_*p*_; the former denotes the amount of signal transported to *ω*, therefore being analogous to *γ*. Let *G*(*s*_*i*_) = *g*(*s*_1_) + … + *g*(*s*_*i*_) be a cumulative distribution function of the noise. Note that 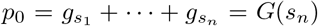. Conceptually, *G* is a different construct than *M* or *N* , since it does not denote the amount of signal present in any given spectrum, but rather the amount of signal *removed* from *μ*.

For centroided spectra, the optimization problem (7) can be rephrased as a minimization over the variables *p* and *g* as follows:

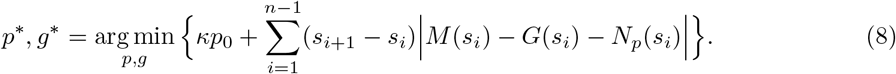

Note that the minimization above is performed over both *p*_*j*_ and *g*(*s*_*i*_) variables. Moreover, we require 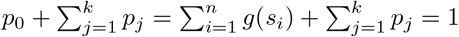, as all the signal in the observed spectrum needs to either be explained by the expected spectra or labeled as noise. The sum of *p_j_* variables denotes the proportion of signal explained by the model spectrum.

Note that the optimization problem (8) is similar to formula (4) from Theorem 2, expressing the Wasserstein distance between a pair of spectra. It is equivalent to computing the distance between the model and the experimental spectrum after removing some of the signal from the latter, and additionally penalizing for the amount of signal removed. However, it does not give any clear physical interpretation of the denoising cost *κ*.

The minimization problem (8) is an example of the Least Absolute Deviation regression [17]. One of the common approaches to solving such problems is by using the technique of linear programming, which optimizes a linear function under a set of linear constraints [18]. However, the function to be minimized in problem (8) is not linear due to the absolute value, and the problem needs to be converted before using linear programming to solve it. There are several ways to convert a Least Absolute Deviation regression problem into a linear program, and choosing the proper one in each particular application is a crucial factor in obtaining a computationally efficient solution. We investigated several approaches and found that, in the case of linear deconvolution of mass spectra, the approach based on the ideas described in [19] seems to be the most efficient.

The derivation of the linear program equivalent to the minimization problem (8) has been relegated to the Appendix. The main idea is to show that solving the deconvolution problem is equivalent to solving the following linear program for the vector of variables *z*, and analyzing the differences between left- and right-hand sides of the inequalities:

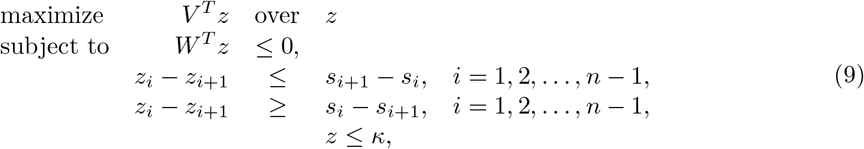

Above, *V* is a vector of the observed spectrum intensities, such that *V*_*i*_ = *μ*(*s*_*i*_), and *W* is a matrix of expected intensities, *W*_*ij*_ = *v*_*j*_(*s*_*i*_). The above problem can be solved using techniques for linear programming such as the Simplex or the Interior Point methods. In our implementation, we have chosen the Simplex method as it can take advantage of the sparsity of matrix *W*.

Note that the linear program above uses only the vectors of intensities and mz values, which are given as the input to the deconvolution problem. Therefore, we do not need to compute any additional values prior to solving the program, such as the cumulative sums used in the Wasserstein distance. Moreover, the *W* matrix tends to be sparse. Modern implementations of the Simplex algorithm can exploit this property to speed up the computations by reducing the number of arithmetic operations.

We have implemented the linear deconvolution method presented above in our Python 3 package, `masserstein`. In the next subsections, we test our implementation on both experimentally obtained and simulated spectra.

### A note about profile spectra

As in the case of computing the Wasserstein distance, the deconvolution algorithm can be applied to profile experimental spectra provided that the sampling of intensity values is uniform over the m/z axis. This is how the results shown in Fig. 1 were obtained, with *κ* = 0.02. According to our knowledge, the Wasserstein distance is the first solution that allows for this kind of processing without the requirement for peak detection or centroiding. In the next section, we validate the performance of this approach on a set of experimentally obtained spectra as well as on simulated data.

## 3 Results

### 3.1 Calibration mix study

In this section we verify whether the deconvolution-denoising method based on the Wasserstein metric can filter out the background noise from experimental spectra, and how accurately it can estimate ion proportions. We also compare the performance of the method on centroid and profile experimental spectra.

As our test dataset, we take 200 repeated measurements of Pierce LTQ Velos ESI Positive Ion Calibration Solution (Thermo Scientific). This calibration mixture is composed of caffeine, a short peptide with the sequence MRFA, and Ultramark 1621, a compound shown in Fig. 3.1. Note that due to varying side group lengths, Ultramark 1621 is in fact a mixture of 13 different compounds. In this dataset, all the isotopic envelopes of the compounds of interest are disjoint. The effect of overlapping isotopic envelopes will be studied in the next subsection.

Due to the different ionization rates of different compounds, it is difficult to predict the correct proportions of ion signals from their concentrations in samples. Therefore, we have assumed that our ground truth to which we compare our estimates are the true signal areas of compounds. To obtain the latter, we have manually selected the signal regions of all compounds, taking into account their monoisotopic peaks and peaks of isotopologues containing one additional neutron (i.e. first isotopic peaks). The region selection was performed on an average spectrum based on the 200 profile spectra. To ensure that the selected regions contain whole peaks in all the spectra, we have additionally taken into account the standard deviation of intensity at each point. The selected regions are shown in Supplementary Table S1, and a fragment of the average spectrum used to select them in Fig. 6. Next, for each spectrum we have integrated the signal within the selected regions using the trapezoidal rule. For each compound, the signals of its monoisotopic and first isotopic peaks were summed to yield the total signal of the compound.

**Figure 6:**
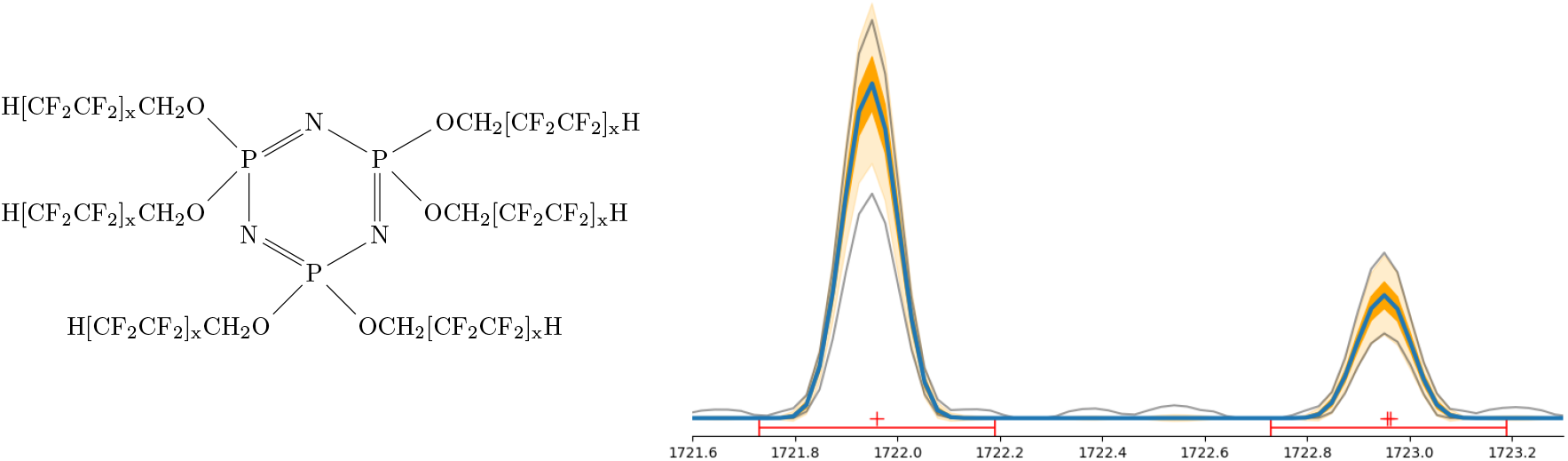
Left: Ultramark 1621, one of the compounds analyzed in this study; x=1,2,3. Right: A fragment of an average spectrum used to define m/z regions of ion signals, showing first two peak of an isotopic envelope of Ultramark 1621 with 14 CF_2_CF_2_ groups. Blue line shows the average signal. Dark orange and light orange ribbons show ±*σ* and ±3*σ* regions, where *σ* is the standard deviation of signal intensity. Grey lines show the maximum and minimum signal over the 200 spectra. Red dots show the m/z values of theoretically predicted peaks. As the theoretically predicted masses agree well with the signal apexes, we infer that the spectrum is properly calibrated. As the maximum and minimum lines are in proximity of the ±3*σ* ribbon, we infer that there are no outlying measurements of intensity. Random increases of maximum signal between the peaks indicate the presence of background noise.

We inspect the results of linear deconvolution for several manually selected values of *κ*. The selection was based on the observed peak widths in profile spectra, which ranged from 0.013 at 195 Da to 0.38 at 1725 Da. In principle, setting *κ* to half base width of the broadest observed peak allows the method to use all the necessary experimental signal. However, we observe that the performance is best for *κ* = 0.4, i.e. the full base width of the broadest peak. This value allows for more flexible transport of signal between peaks of an isotopic envelope, effectively countering the effect of the variance of experimental peak intensity. The effect is mostly pronounced for low intensity peaks with small signal to noise ratio, in which case the particularly high variance of signal intensity has a detrimental effect on the estimation unless *κ* is sufficiently high. The reason behind this effect is explained in more detail in the following paragraphs.

#### Analysis of centroided spectra

The first goal of this study is to verify whether the linear deconvolution algorithm implemented in the `masserstein` package returns accurate results when applied to a centroided experimental spectrum. In order to perform the centroiding in a controlled manner, we have implemented our own peak-picking procedure. Briefly, the spectra are centroided by integrating the signals within regions delimited by 0.2 of the apex intensity. The pseudo-code of the full procedure is shown in Algorithm 3 in section 4.4. We have validated our implementation by comparing the centroided peak intensities with the manually integrated signal areas and found a good agreement.

We have generated the theoretical spectra of the compounds of interest using IsoSpecPy [20], assuming a hydrogen adduct. In each spectrum we have retained the monoisotopic and the first isotopic peaks, so that the theoretical spectra correspond to the manually identified regions. After that, we have used `masserstein` to deconvolve all of the 200 experimental spectra with respect to the theoretical spectra. We have inspected several values of the denoising penalty *κ* ranging from 0.05 to 0.6. For *κ* = 0.4, the deconvolution of all spectra took around 98 seconds on a single core of an Intel(R) Core(TM) i5-6300HQ CPU @ 2.30GHz processor

The `masserstein` package requires that all the spectra are normalized. Prior to normalization we have computed the total ion currents of the experimental spectra. After obtaining the estimated proportions of ions, they were multiplied by the total ion current of the experimental spectrum. This way we obtain the total signal explained by each ion, which we then compare to the manually calculated signal area.

The results for *κ* = 0.4 are shown in Fig. 7. The overall correlation between the estimated and the manually computed signal was equal to *ρ* = 0.9998. The mean difference between the estimated and manually integrated signal relative to the latter was equal to 1%, indicating a slight downward bias for this value of denoising penalty. The mean absolute relative difference was equal to 0.017, meaning that the average error of the estimation is equal to 1.7% of the true value. The detailed results for each ion are shown in Supplementary Fig. S6.

**Figure 7:**
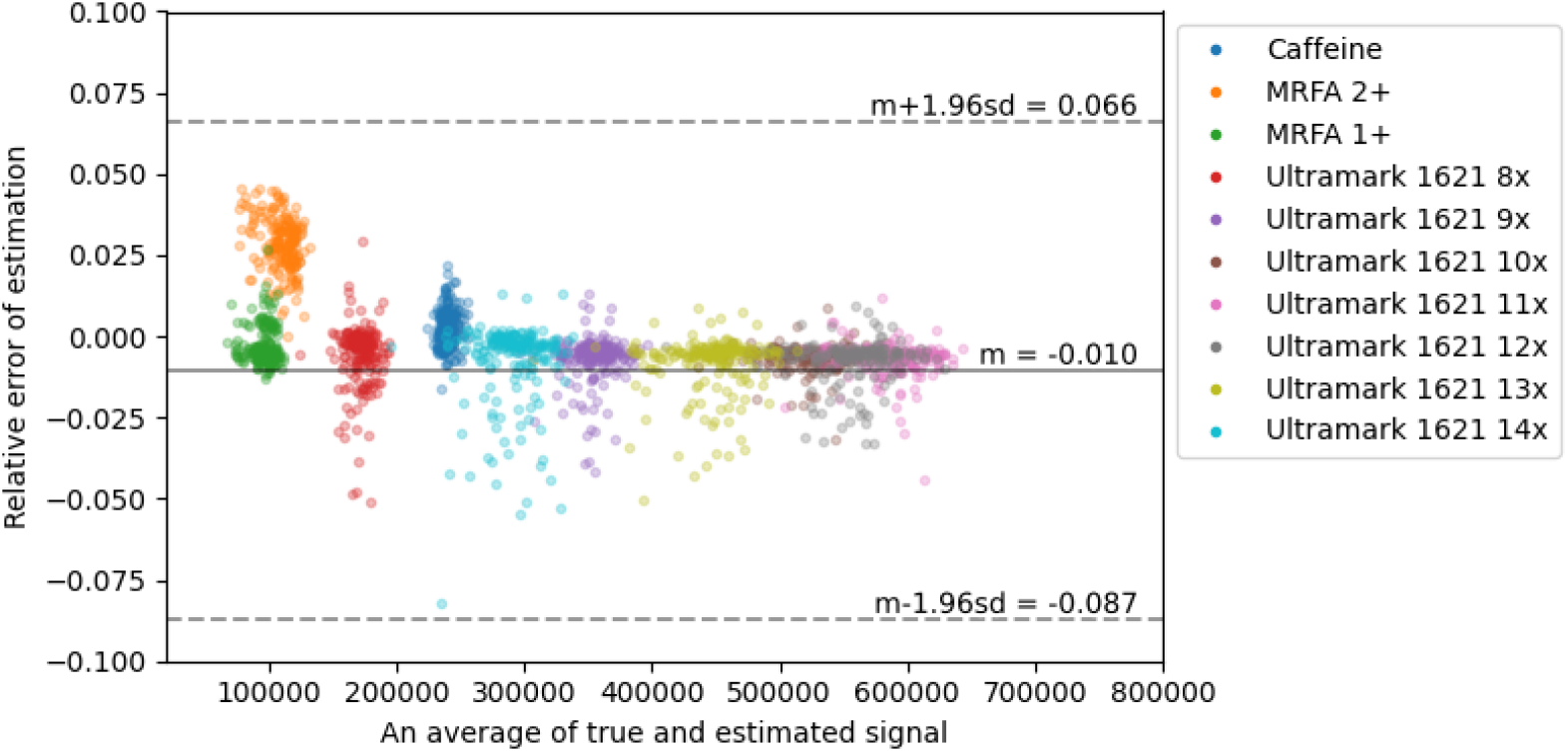
A Bland-Altman plot summarizing the deconvolution results for the calibration mix on centroided spectra. Each point corresponds to an estimate of an ion intensity in one of the 200 spectra analyzed, with colors corresponding to different ions. The Y coordinate of each point corresponds to the estimated minus the true signal divided by the latter. Ultramark 1621 8x denotes Ultramark 1621 with 8 CF_2_CF_2_ groups, etc.

For *κ* = 0.3 the results were similar for most ions, and the overall correlation of the estimated and true signals was equal to *ρ* = 0.9994. However, we have found a strong down-estimation of the signal of Ultramark 1621 with 15 CF_2_CF_2_ side groups, which caused the drop in the correlation. We have found out that the bias is caused by a large variability of the relative peak intensities of this ion. In some spectra, the first isotopic peak was up to two times lower than on average. Such a large variability is most likely caused by a small number of ions of this compound.

Highly variable peak heights, combined with low denoising penalties, are detrimental to our current implementation of `masserstein`. When the maximum feasible transport distance is low, the procedure necessarily fits to the smallest matching peak of the experimental spectrum. This is because after all the signal of such peak is transported to a theoretical spectrum, there is no neighboring signal left that can be feasibly transported.

Increasing the denoising penalty allows to distribute the experimental signal more evenly over the theoretical spectrum, therefore increasing the accuracy of the estimation. However, when the penalty is too high, the background noise may also be transported to the theoretical spectrum, leading to an overestimation of an ion’s signal.

The results demonstrate that the optimal transport theory applied to the deconvolution problem is capable of giving very accurate estimates of ion signals, provided that all the considered signals are over the limit of quantification to ensure small variability of peak areas. As we show below, it also opens the possibility of directly analyzing the profile spectra without the peak-picking step.

#### Analysis of profile spectra

In the current implementation of the deconvolution algorithm, we treat profile spectra in the same way as the centroided ones. As discussed in the previous section, this approach gives a good approximation when the signal sampling is uniform over the m/z axis. However, this is often not the case for spectrometers which have a non-constant resolving power. As peaks get broader with the increasing m/z value, less data points are needed to reflect the signal shape. This phenomenon is often exploited in order to decrease the data size. One of the way to circumvent this problem in order to use the current implementation of our method is to resample the signal intensities.

We have resampled our spectra using a piecewise linear interpolation, in which the signal intensity in each point is approximated by a weighted average of the neighbouring intensities. The full procedure is shown in pseudo-code in Algorithm 2 in section 4.4 of the Appendix. For each spectrum, we have interpolated its signal such that the spacing between neighboring m/z values was 0.001.

The downside of the resampling strategy is a large increase of the data complexity, and consequently, the computational time. For the deconvolution of the 200 profile spectra with *κ* = 0.4, the computational time increased from 98 seconds to 35 minutes.

The penalty *κ* = 0.4 yielded an overall correlation of 0.9998. The results for all compounds are shown jointly in Fig. 7, and for each ion in detail in Supplementary Fig. S7. We have noticed a systematic slight overestimation of the caffeine signal, most likely due to incorporation of a small unidentified peak at 195.72 Da which was present in most spectra. For *κ* = 0.3 we have observed a bias in the estimation of Ultramark 1621 with 15 CF_2_CF_2_ groups, similar as for the centroided spectra.

In general, the results were similar to the ones obtained on centroided spectra. This shows that `masserstein` allows for the processing of profile spectra without the need of peak peaking. Further research into applications of the optimal transport theory to the processing of mass spectra has the potential to increase both the computational efficiency and the accuracy of the results.

#### Overlapping isotopic envelopes

In our final experiment on the calibration mix data, we investigate the influence of overlapping envelopes on the accuracy of the estimation. In the case of disjoint isotopic envelopes, like in the previous experiments, it is easy to obtain the ground truth by manually selecting peak regions for integration. However, the task is complicated when the peaks of the envelopes overlap, because manual integration does not allow to separate them and compute their individual signals. Therefore, we have decided to simulate this effect using the calibration mix data.

For each spectrum in profile mode, we have created its copy shifted by one hydrogen mass. Each spectrum was mixed with its shifted copy in proportion 0.7 of the original and 0.3 of the copy. The spectra were subsequently centroided as in the previous examples.

To generate a model spectrum, we have taken the formulas from the previous experiments, and the same formulas with one additional hydrogen. When adding a hydrogen atom to the sum formula, we only modified the ion’s mass, while keeping its charge unchanged. Note that, from a computational perspective, it does not matter whether the formulas obtained this way correspond to any actual chemical compound.

From the computational point of view, this dataset is much more difficult than the previous ones, and we expect a decrease in estimation accuracy. One of the reasons is that when overlapping signals merge, the apex positions shift relative to the original ones. Therefore, peak picking a profile spectrum with overlapping signals returns distorted peak positions, and for a sufficiently large overlap one gets a single peak instead of two. This poses a difficulty for approaches based on pointwise comparison of spectra, while the Wasserstein distance is more robust to such changes.

Using `masserstein`, we have deconvolved the mixed spectra with denoising penalty *κ* = 0.4. This time, the deconvolution of all the spectra took approximately 3 minutes (not including the time needed for centroiding). The results were compared with the signal areas integrated in the previous experiment, rescaled either by 0.7 or 0.3 to accommodate for the mixing proportions. The correlation between the estimates of `masserstein` and the true signals was equal to *ρ* = 0.9985, only slightly smaller than for the previous datasets. The detailed results are shown in Supplementary Fig. S8.

Additionally, we have calculated the ratios of estimated proportions of corresponding ions. In each spectrum we have compared the estimated signal of an original ion to the estimated signal of it’s counterpart with one additional hydrogen atom. We have compared the ratio obtained this way to the reference value of 0.7/0.3 ≈ 2.33. The result is shown in Fig. 8.

**Figure 8:**
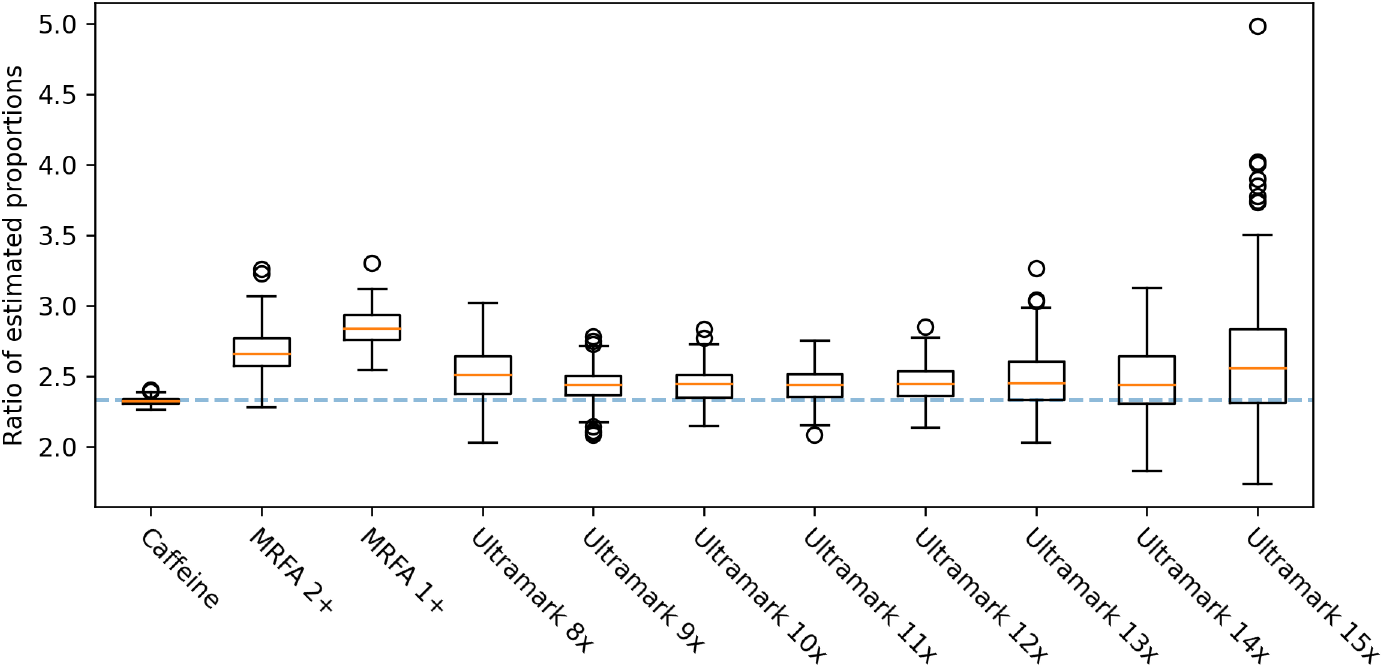
The ratio of estimated signals of ions with overlapping isotopic envelopes. Each boxplot corresponds to a pair of ions with sum formulas differing by one hydrogen atom. The dashed line represents the true ratio.

We have observed that the ratio is overestimated for MRFA 1+. Comparing the ratio with the results shown in Supplementary Fig. S8, we conclude that this is caused by an overestimation of the signal of the original MRFA 1+ ion (i.e. without the added hydrogen atom). For all the other ions, the true ratio was within the 95% confidence interval of the estimation. The estimated ratio showed a high variance for Ultramark 15x, which is likely caused by the lower signal to noise ratio of this ion compared to the other ions. On the other hand, the estimated signal ratio was the most accurate for caffeine, likely due to low amount of background noise in the neighbourhood of its isotopic envelope and a sufficiently high signal intensity.

The results of the three experiments presented in this subsection show that our deconvolution algorithm implemented in `masserstein` is capable of accurate estimation of signal intensity based on experimental spectra in both centroided and profile mode, and also in the presence of overlapping isotopic envelopes. However, the dataset presented in this section contained only a handful of molecular formulas. In order to verify the results for a broader range of molecules, and to demonstrate that `masserstein` is not limited to any particular type of ions (be it lipids, peptides or metabolites), in the next subsection we perform an extensive analysis on simulated datasets consisting of purely random molecular formulas.

### 3.2 Computational experiments

In this section, we evaluate the accuracy and bias of the estimation performed by `masserstein` using simulated mass spectra. We introduce a series of measurement distortions into the observed spectrum in order to reflect the variability of peak heights and limited resolving power and accuracy observed in real spectra.

We have created several simulated datasets by computing spectra of mixtures of randomly generated molecules. In each dataset, all the molecules had the same nominal mass in order to ensure high overlap of their isotopic envelopes. The molecules were simulated by subsequently sampling elements in the order of C, O, N, S, and P. Note that any user-supplied formula can be used in `masserstein`, as no mathematical procedure used in this work depends on the chemical properties of molecules. Therefore, in order to simplify the simulation procedure, we did not restrict the sampled molecules to ones which are chemically possible.

We have simulated datasets consisting of isobaric molecular formulas for a range of nominal masses from 60 to 12 000 Da and from 1 to 8 isotopic envelopes. Based on the formulas, we have generated theoretical spectra using IsoSpecPy [20]. To obtain simulated experimental spectra, for each dataset we have mixed the theoretical ones in random proportions. We have generated both centroid and profile experimental spectra. In the latter case, we have assumed a Gaussian shape of peaks.

To each experimental spectrum, in both profile and centroid mode, we have introduced extensive distortions to simulate the effects of a finite number of molecules, electronic noise, measurement inaccuracy in mass and intensity domain and limited resolving power (100 000 at 600 Da in the case of profile spectra). The procedure is described in detail in section 4.3 of the Appendix.

To assess the quality of deconvolution results, we calculate the mean absolute deviation (MAD) between true and estimated signal contributions of the theoretical spectra:

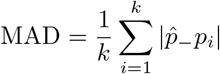

In the above formula, *k* is the number of theoretical isotopic envelopes, *p*_*i*_ is the true proportion of the *i*-th envelope, and 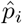 is its estimated proportion. We do not directly compare the amount of signal that our method classifies as noise, *p*_0_, as this information is implicitly included in the sum of the estimated proportions.

In the case of profile observed spectra, we have inspected two denoising penalties, equal 0.0075 (half peak base width) and 0.08. The results are shown in Supplementary Fig. S10. For the lower penalty, the MAD was mostly between 10^−2^ and 10^−1^, indicating at least one accurate decimal digit in an average estimate. We have observed some bias in the estimation, as the estimated proportions were usually lower than the true ones. The mean of the residues was equal to −0.0197, while their standard deviation was 0.025. The bias increased with increasing true proportion. This can be explained by the fact that the isotopic envelopes of abundant ions have a larger variance of their peak heights due to isotopologue sampling. For the 0.08 penalty, the MAD was smaller and usually around 10^−2^. The decrease in MAD was especially pronounced for low numbers of isotopic envelopes. One of the reasons for better performance in this case is a lower bias, as the mean of the residues was −0.008. However, at the same time, their variance increased to 0.028. In all cases, the processing of a single spectrum took less than one second on a laptop computer with Intel(R) Core(TM) i5-6300HQ CPU @ 2.30GHz processor.

In the case of centroided spectra, the denoising penalty was set to 0.02 (ten times the standard deviation of the simulated m/z measurement error). The results are shown in Supplementary Fig. S9. The MAD was mostly between 10^−3^ to 10^−2^, indicating at least two accurate decimal digits an average estimation. There were no clear differences between low and large masses, indicating that isotopic envelopes of 200 Da ions have enough information for accurate deconvolution using our method. In most cases, the best estimates per observed spectrum had at least three accurate decimal digits. We did not observe any bias of the estimation; the mean of the residues was one order of magnitude lower than in the case of profile spectra.

The results show that our method is able to perform accurate estimation of molecule proportions even in the case of several isobaric interferences and additional chemical noise. Moreover, it is applicable to both profile and centroided spectra. In the case of profile spectra, we have observed a tradeoff between the estimation bias and variance for different values of the denoising penalty *κ*. The results presented above were better for centroided spectra. However, the two different types of distortions used to simulate both types of observed spectra are not directly comparable, and their magnitudes differ.

Notably, we have obtained accurate results for up to 6 isobaric molecules of 200 Da in the presence of 50 additional interfering peaks, even though the molecules themselves have only a few peaks in their isotopic envelopes. In this case, the largest absolute error of estimation per centroided spectrum, averaged over 100 replicates, was 0.026 (see Supplementary Fig. S9). This indicates that, on average, all the estimates had at least one accurate decimal digit.

## 4 Conclusions and discussion

In this work, we present advances in theoretical studies on the problem of linear deconvolution of mass spectra, defined as fitting a linear combination of theoretical spectra to an experimental one. This problem is ubiquitous in mass spectrometry, appearing either implicitly or explicitly in areas as diverse as metabolomics, proteomics and polymer science. Furthermore, the theoretical foundations of the methods studied in this work are not limited to mass spectra, and can be readily applied to areas such as nuclear magnetic resonance spectroscopy as long as reference spectra are available. The broad range of applications is achieved by focusing on an abstract problem of fitting a linear combination of reference signals to an experimentally measured one, as opposed to developing a method restricted to a particular application.

One of the major factors that hinder the performance of currently available linear deconvolution methods is the fact that the locations of experimentally measured peaks never match the theoretically predicted ones perfectly. This is caused, among others, by measurement inaccuracies, which, even if seemingly small, are unavoidable. Although they may be negligible when the spectra are analyzed manually, an m/z difference as small as a millionth of a Dalton means that a computer program sees the peak locations as different.

In order to circumvent this problem, the currently available software matches peaks within so called mass windows of predefined width [4, 1]. The width needs to be specified by the user, which makes it more difficult to apply this kind of approach in practice. The choice of a mass window width is further complicated by the occurrence of highly overlapping peaks in profile spectra. In some cases, the individual apexes of such peaks are no longer visible. Instead, we obtain a single apex located between the “true” apexes of the component peaks. This kind of merging of peaks, occurring especially in complex profile spectra, leads to an increased difference between theoretical and observed peak locations in centroided spectra. Therefore, especially in complex or low-resolution spectra, the optimal window width may be considerably greater than the nominal accuracy of the instrument.

Even if the user knows the optimal width of a mass window, this kind of approach has several intrinsic drawbacks. Due to the infinite resolution of a theoretically predicted spectrum, a mass window usually contains several theoretical peaks. Within the window, those peaks are effectively treated as one. Therefore, this procedure effectively limits the resolution of the *in silico* predicted spectrum, leading to an unnecessary loss of information. On the other hand, it tends to merge closely positioned signal and noise peaks in the experimental spectrum, which influences the estimated proportions.

In order to alleviate those difficulties, we have investigated the application of the Wasserstein distance to the problem of linear deconvolution. Methods based on quantifying the distance in the m/z domain needed to transform one spectrum into the other are naturally robust to limited resolving power and accuracy of instruments. This robustness makes them a promising tool for methods based on comparing experimentally acquired spectra.

The pre-existing, extensive mathematical research on the topic of optimal transport has resulted in powerful theorems that allow to express the Wasserstein distance as a computationally feasible integral. Those theorems have allowed us, in turn, to develop an effective algorithm for linear deconvolution that does not require any optimization of the transport plan. Instead, the optimal cost of transporting the signal between spectra is computed in a straightforward manner using the aforementioned theorems, and a technique of linear programming is used to directly optimize the proportions of theoretical spectra.

### The choice of the denoising penalty

Our method requires the user to specify a single parameter: the cost of denoising *κ*, interpreted as the maximum feasible transport distance in the m/z domain. This interpretation of *κ* allows to treat it analogously to the radius of a mass window. However, a major difference between the two approaches is that *κ* acts as a *soft threshold*: during deconvolution, whether some signal is transported between two peaks depends not only on their distance relative to *κ*, but also on the shape of the theoretical isotopic envelopes. This allows for some imprecision in setting the value of *κ* and makes the results more stable when the value is sub-optimal.

Similarly to the width of a mass window, in practical applications the choice of *κ* is not straight-forward. The performance of the deconvolution methods for given parameter values is influenced by several factors, including the instrument accuracy and resolving power. Usually, the results for several different values need to be inspected manually in order to make a final decision. The half-base widths of peaks in profile spectra and the instrument accuracy serve as a convenient reference for the reasonable range of values of this parameters. Setting *κ* higher than 1 is usually inadvisable, as it allows to transport the signal between peaks of ions with different atomic compositions.

The choice of *κ* resembles the classical variance-bias trade-off known from the field of machine learning. Small values of *κ* lead to down-estimation of ion proportions, as insufficient signal is available to be transported onto their theoretical isotopic envelopes. On the other hand, while high values of *κ* allow to transport all the required signal, they lead to incorporation of noise in the estimated proportions, therefore increasing the variance of the estimation. Additionally, if systematic sample impurities are present, high values of *κ* lead to over-estimation. Further studies are needed in order to give precise guidance as to the optimal value of this parameter.

### Further research

The Wasserstein distance quantifies the similarity of spectra only in terms of the difference in m/z locations of signals. A drawback of this approach is that it is relatively sensitive to differences in peak intensities. This is further emphasized by the different nature of measurement uncertainties in the mass and the intensity domain, caused by the fact that mass spectrometers measure the m/z and the number of ions differently. Although accounting for this phenomenon would be desirable, it is highly non-trivial to formalize it mathematically in a way that would be suitable for applications in mass spectrometry and linear deconvolution in particular. Especially in the latter, allowing peak intensities to vary may lead to instability of the results—when everything is variable, we can fit anything to anything. For this reason, in this work we only allow the signal to be removed from the experimental spectra, and not from the theoretical ones.

A practical consequence of allowing the signal to only be removed from the experimental spectrum is that, when simulating the theoretical spectra, we need to discard peaks that are under the level of quantification. The measured intensities of such peaks are unreliable, leading to erroneous results for the whole ion. In an extreme case, when a peak is missing in the experimental spectrum but is present in the theoretical one, it forces the estimated proportion to be zero. The lack of experimental intensity that can be transported onto this theoretical peak means that the whole isotopic envelope needs to be discarded. Allowing for some flexibility in the intensities of theoretical peaks would allow to alleviate this difficulty. However, in order to accurately reflect the observed variability of peak intensities, the deconvolution procedure needs to be coupled with a mathematical model of the shape of experimentally measured isotopic envelopes. Whether such coupling is mathematically and computationally feasible remains an open question.

### Implementation

We have implemented the discussed algorithms in a Python 3 package called `masserstein`. Our implementation is designed to be applicable in larger data processing pipelines. Efficient development of pipelines requires freely available modular tools, which perform specific tasks and can be easily combined. Therefore, our implementation does not perform any additional pre- or postprocessing of the results, such as peak-picking or correcting for proton affinity of molecules. Such procedures can be performed separately using designated tools, available e.g. in the OpenMS package [21].

On a final note, we reiterate that, from a methodological point of view, molecule identification and quantification are two separate tasks. Accordingly, in this work, we assume that the chemical formulas of the molecules to be quantified are known a priori. These may come either from the scientific question at hand (such as the frequency of a given posttranslational protein modification), from the knowledge about the experimental setup (such as whether lipid extraction was performed), or from an identification study performed prior to quantitative analysis (e.g. using the SIRIUS program [22]).

## Acknowledgements

This work has been supported by the Special Research Fund (BOF) of the Hasselt University BOF18BL09, the FWO-PAS grant VS.028.19N, and the Polish National Science Centre grants #2018/29/B/ST6/00681 and #2017/26/D/ST6/00304.

## Appendix

## 4.1 Derivation of the denoising procedure

In the main body of the article, we have presented two formulations of an optimization problem that allows for simultaneous spectrum denoising and estimation of ion proportions (Eqs. (7), (8)):

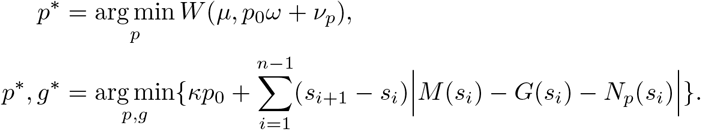

In this section, we show a proof of the equivalence of the formulas. We also give further examples which illustrate several properties of our denoising-deconovolution procedure, such as the relationship between the denoising penalty *κ* and the maximum transport distance. Those examples further motivate the interpretation of *κ* as a *soft threshold* on the transport distance.

Consider a set of theoretical spectra *v*_*j*_ and an observed spectrum *μ*. Recall that the latter is assumed to be composed of theoretical spectra and some additional signal, referred to as the chemical noise. Formally, we assume that there are some *true proportions* of spectra *v*_*j*_, denoted *p*_*j*_, satisfying

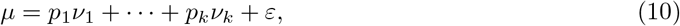

where *ε* is a spectrum representing the chemical noise. We assume that *p*_1_ + … + *p*_*k*_ ≤ 1, allowing some of them to be zero. Furthermore, recall that we assume that all the spectra *μ* and *v*_*j*_ are normalized by their total ion current. It follows that the total signal intensity in *ε* is equal to 1 − *p*_1_ − *p*_2_ − … − *p*_*k*_, which will be denoted as *p*_0_.

Under the assumption that *ε* is empty, the optimal proportions 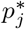 were found by minimizing the Wasserstein distance *W*(*μ, p*_1_*v*_1_ + … + *p*_*k*_*v*_*k*_) between *μ* and a linear combination of *v*_*j*_ [12]. With non-empty *ε*, we need to incorporate it’s removal to the estimation procedure. In order to do that, we introduce an auxiliary spectrum *ω*, and model the denoising as transporting signal from *μ* to *ω*. That is, the amount of signal transported from *μ*(*s*_*i*_) onto *ω* is interpreted as the amount of noise signal at *s*_*i*_.

The auxiliary spectrum *ω* may be a somewhat non-intuitive concept. First, we assume its total signal sums up to one, but we do not explicitly assume this signal to have any particular m/z value. Second, we assume that there is a constant cost of transporting signal from *μ* to *ω*, denoted as *κ* and referred to as the *denoising penalty*. In a way, *ω* can be thought of as being equidistant to all signals in *μ*. The reason for defining *ω* in such way is to formally define a cost of denoising that does not depend on the m/z value of the removed signal.

Now, we look for optimal proportions as

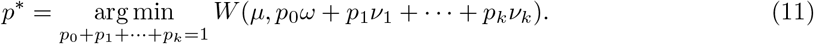

That is, we look for proportions that allow for the optimal transport of the signal from *μ* onto *v*_*j*_’s and *ω*.

The Wasserstein distance between two measures *μ* and *v* defined on a space 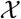 is given by

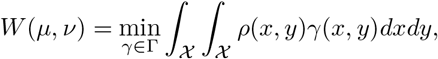

where Γ is the space of all joint distributions of *μ* and *v* and *ρ*(*x, y*) is a distance function between points *x* ∈ *X* and *y* ∈ *Y*. In general, the definition works for any distance function *ρ*. In our case, we define *ρ*(*x, y*) between two m/z values *x* and *y* as follows:

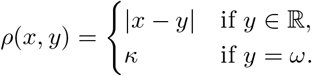

Based on the above definitions, we have

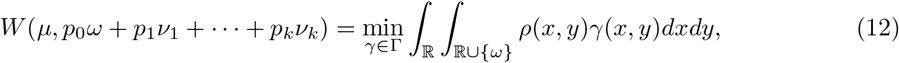

where the minimization is over all joint distributions *γ* of *μ* and *p*_0_*ω* + *p*_1_*v*_1_ + … + *p*_*k*_*v*_*k*_. It follows that *γ* satisfies the following properties:

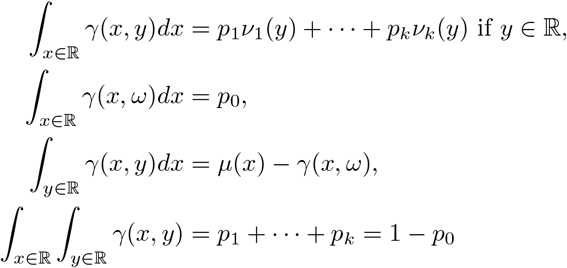

We now proceed to convert the optimization problem (12) to a form that is computationally feasible. We start by splitting the integral over ℝ ∪ *ω* into two summands and simplifying the resulting terms:

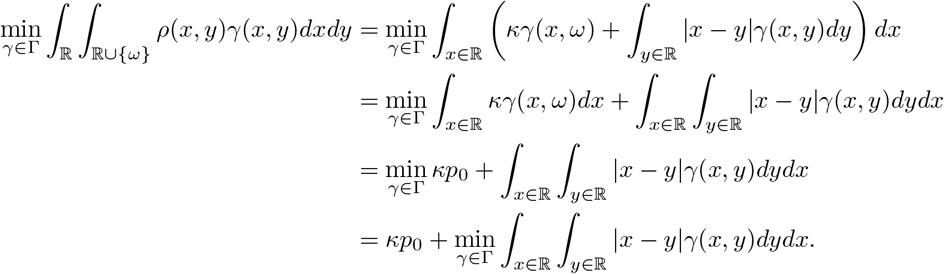

In the last line, we arrive at the total cost inflicted by signal removal, *p*_0_*κ*, and a double integral that is strikingly similar to the definition of the Wasserstein distance between two spectra. Namely, we integrate the transport distance |*x* − *y*| multiplied by *γ*(*x, y*), the amount of signal transported between *x* and *y*. However, unlike in the definition of the Wasserstein distance, now Γ is a set of joint distributions over ℝ and ℝ ∪ *ω*, so *γ* function may not be a joint distribution of two probabilistic measures defined on the real line. It means that we cannot yet use the formula that joins the Wasserstein distance to the cumulative distribution functions of the compared measures.

To circumvent the above problem, we proceed as follows. Define a measure *g*(*x*) = *γ*(*x, ω*) and denote it’s cumulative distribution function as *G*(*t*). Observe that the total signal in *g* is equal to *p*_0_. It follows that

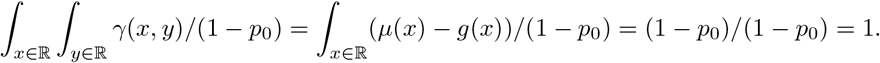

We write 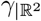 to explicitly denote *γ* function restricted to ℝ^2^. From the above integrals it follows that 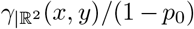 is a two-dimentional probabilistic measure on ℝ^2^. Its marginal measures are

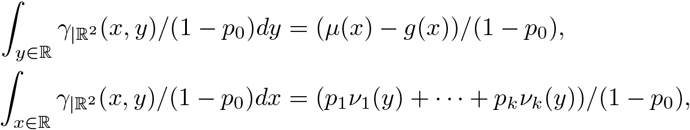

which are both probabilistic measures on ℝ. Note that in both equations above we consider 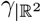, meaning that we assume *y* ≠ *ω*. We now can use Theorem 2 which expresses the Wasserstein distance between two centroided spectra, *μ* and *v*, in terms of their cumulative distribution functions, *M* and *N*. In the first step, we split the minimization over Γ into two steps: first, minimization of 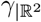, and then minimization of *g*.

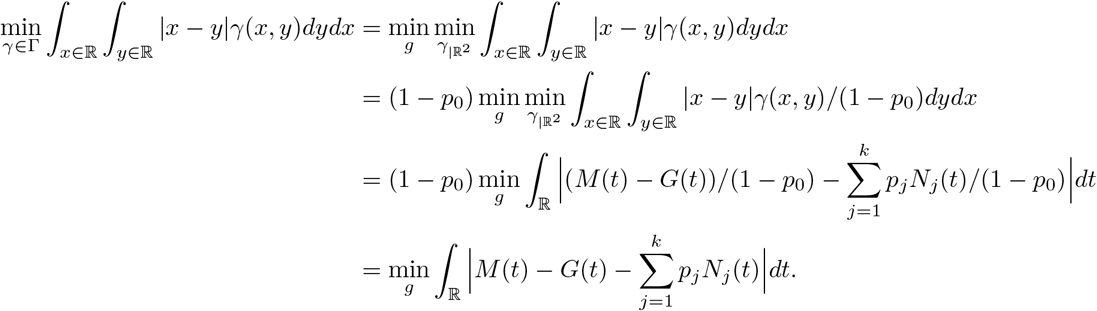

Putting it all together, we arrive at

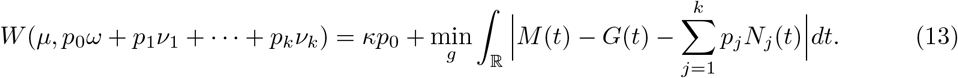

The *κp*_0_ term in the above equation is the penalty for removing *p*_0_ of the signal from the observed spectrum. The minimized integral is the Wasserstein distance between the denoised observed spectrum and the combination of expected spectra under the optimal denoising plan *g*, on the condition that we remove *p*_0_ of the signal.

To obtain the optimal proportions, we minimize the equation over the proportions *p*. Since the term *κp*_0_ does not depend on the denoising plan *g*, as long as *p*_0_ of the signal is removed, we can minimize over both *p* and *g* together and write the following formula for optimal proportions:

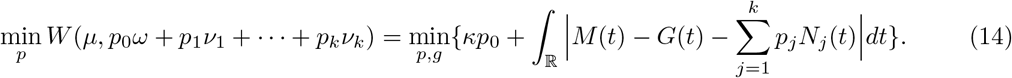

Formally, we minimize over all proportions *p*_0_ to *p*_*k*_ and over the denoising plan such that *g_i_* sum up to *p*_0_. This is however equivalent to minimization over *p*_1_ to *p*_*k*_ (i.e. without *p*_0_) and *g*_*i*_ such that 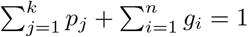. This formulation of the minimization problem is used in the next section.

Before we proceed to describe the algorithm for solving the minimization problem, we give some additional remarks about the problem itself. First, the denoising penalty *κ* is interpreted as the distance between any peak from the observed spectrum to the auxiliary spectrum *ω*. If, for a given observed peak, all the theoretical peaks are further away than *κ*, then *ω* is the closest spectrum to this observed peak. Therefore, *κ* can be interpreted as the maximum ‘feasible’ transport distance. This interpretation is helpful in estimating reasonable denoising penalties. However, it should be treated as an intuition or a rule of thumb rather than a formal property, as transport for distances greater than *κ* might occur in some cases.

An example where a long distance transport is beneficial is shown in Fig. S1. In this example, we “deconvolve” an experimental spectrum with respect to one theoretical spectrum. Both spectra are abstract examples which serve for a simple illustration of the properties of our method, and do not correspond to any actual molecule.

The theoretical spectrum is composed of three peaks with m/z values 1, 2 and 3 Da, and intensities 1/2, 1/2 − *h, h* with 0 < *h* < 1/2. The experimental spectrum is identical to the theoretical one, except that the peak at 3 Da is shifted 1 Da to the right. We analyze two limiting scenarios that may occur in this situation: either the shifted peak is removed, or it’s not.

In the first scenario, no denoising is performed, and therefore the proportion of the theoretical spectrum is equal to 1 and the shifted peak is transported onto its theoretical counterpart. The Wasserstein distance in this case is therefore equal to the height of the shifted peak, denoted *h*.

In the second scenario, the shifted peak is removed. The proportion of the theoretical spectrum is equal to the amount of the remaining experimental signal, that is 1 − *h*. In order to compute the cost of the signal transport in this case, we remove the shifted peak from the experimental spectrum, multiply the theoretical peak intensities by 1 − *h*, and compute the cost of the optimal transport between the resulting spectra.

The CDF of the experimental spectrum after this procedure becomes equal to

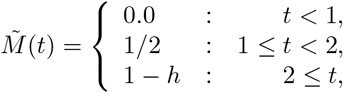

and the one of the theoretical spectrum becomes equal to

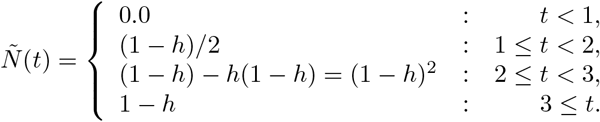

The cost of signal transport between the two spectra is therefore equal to

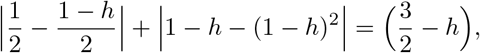

and the total cost of this scenario, obtained by adding the cost of the peak removal to the above cost of transport, is equal to *hκ* + *h*(3/2 − *h*) = *h*(*κ* + 3/2 − *h*), and is higher than the cost of the first scenario whenever *h* < *κ* + 1/2. However, since 0 < *h* < 1/2 and *κ* ≥ 0, the first scenario is always less costly than the second one. Therefore, regardless of the value of the denoising penalty, it is always beneficial to transport the rightmost experimental peak to its theoretical counterpart rather than remove it.

The phenomenon described above occurs because removing a peak induced large disturbance in the optimal transport plan, depicted in Fig. S1. In the first of the described scenarios, the peaks are matched one to one, and only the rightmost experimental peak needs to be transported. On the other hand, in the second scenario, some portion of the signal needs to be transported from each experimental peak, because their intensities exceed the ones of their theoretical counterparts.

In general, *κ* sets a threshold on the maximum transport distance for those peaks of the experimental spectrum which can be removed without causing major distortions in the optimal transport plan or the optimal proportions of theoretical spectra. An example of such case is shown in Fig. S2. In this example, *κ* does define a strict limit on the maximum transport distance. The computation of the costs of the scenarios is done in the same way as in the previous examples.

Another property of our method that should be noted is that the solution to the minimization problem may not be unique in some cases (see Fig. S3 for an example). In the current implementation, we do not attempt to make it unique. Instead, when there are several equally good solutions, we simply pick one at random.

## 4.2 Finding the optimal proportions

In the previous section, we have derived the formula that needs to be minimized in order to obtain optimal proportions of the theoretical spectra within the observed spectrum,

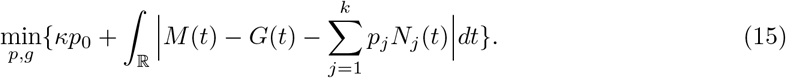

In this section, we show a computational procedure of finding the proportions *p*_*j*_ and the amounts noise *g*(*s*_*i*_). In principle, the above formula could be used to treat profile and centroided spectra differently. The cumulative distribution functions of the profile spectra are continuous, while the ones of centroided spectra are step functions. However, in the current implementation we treat both types of spectra in the same manner. A profile spectrum is treated simply as a particularly long peak list, or a series of intensity measurements at discrete m/z values. It follows that all our CDFs are step functions.

**Figure S1:**
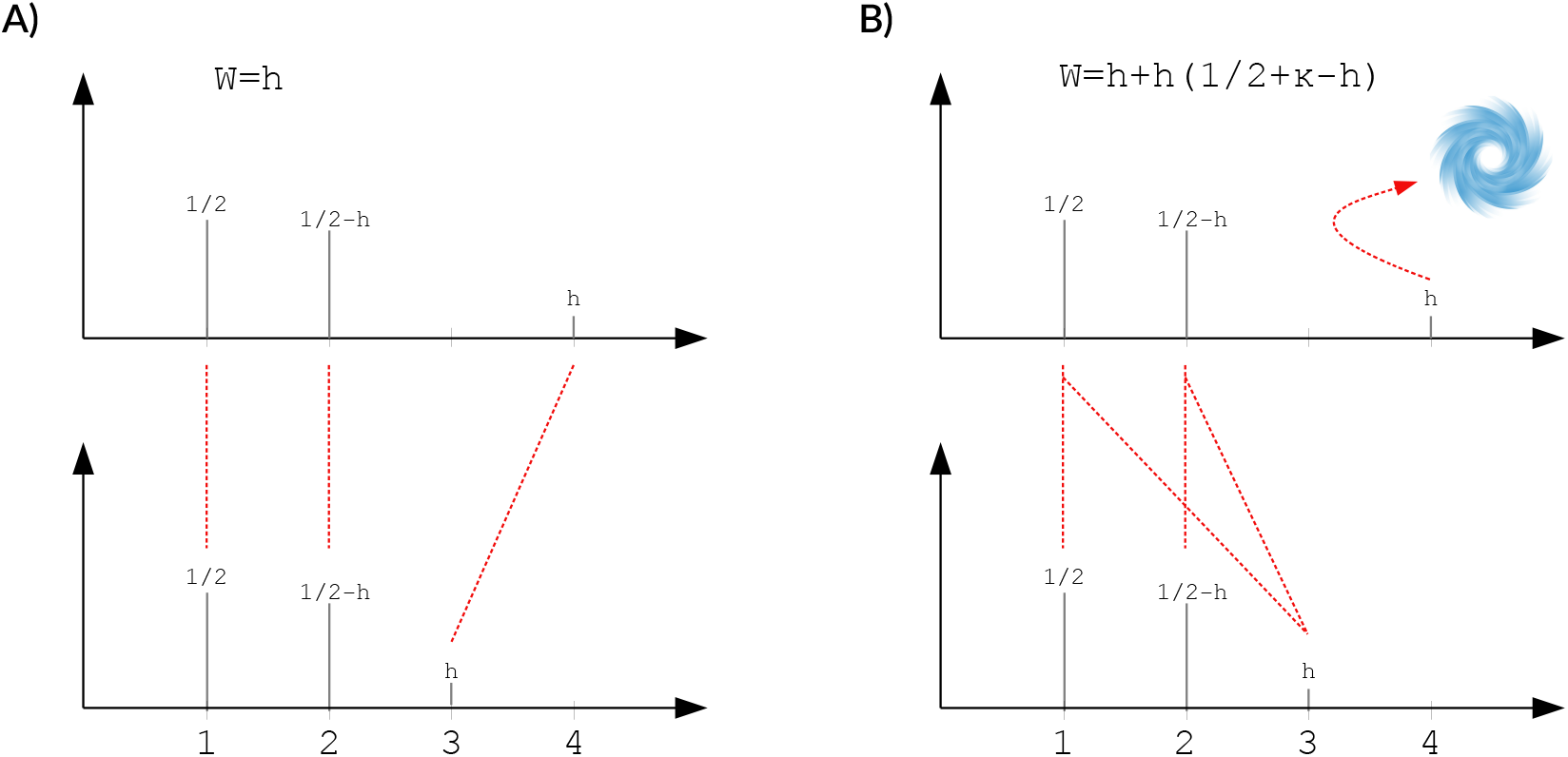
An example of a long-distance transport. The observed spectrum is shown at the top, the theoretical one at the bottom. The spectrum *ω* is represented as a vortex. Two transport plans are compared: A, no denoising, and B, one removed peak. Scenario A is less costly than B regardless of the denoising penalty, and in fact optimal in this example. Long distance transport occurs, because removing the rightmost peak from the experimental spectrum highly disturbs the transport plan.

**Figure S2:**
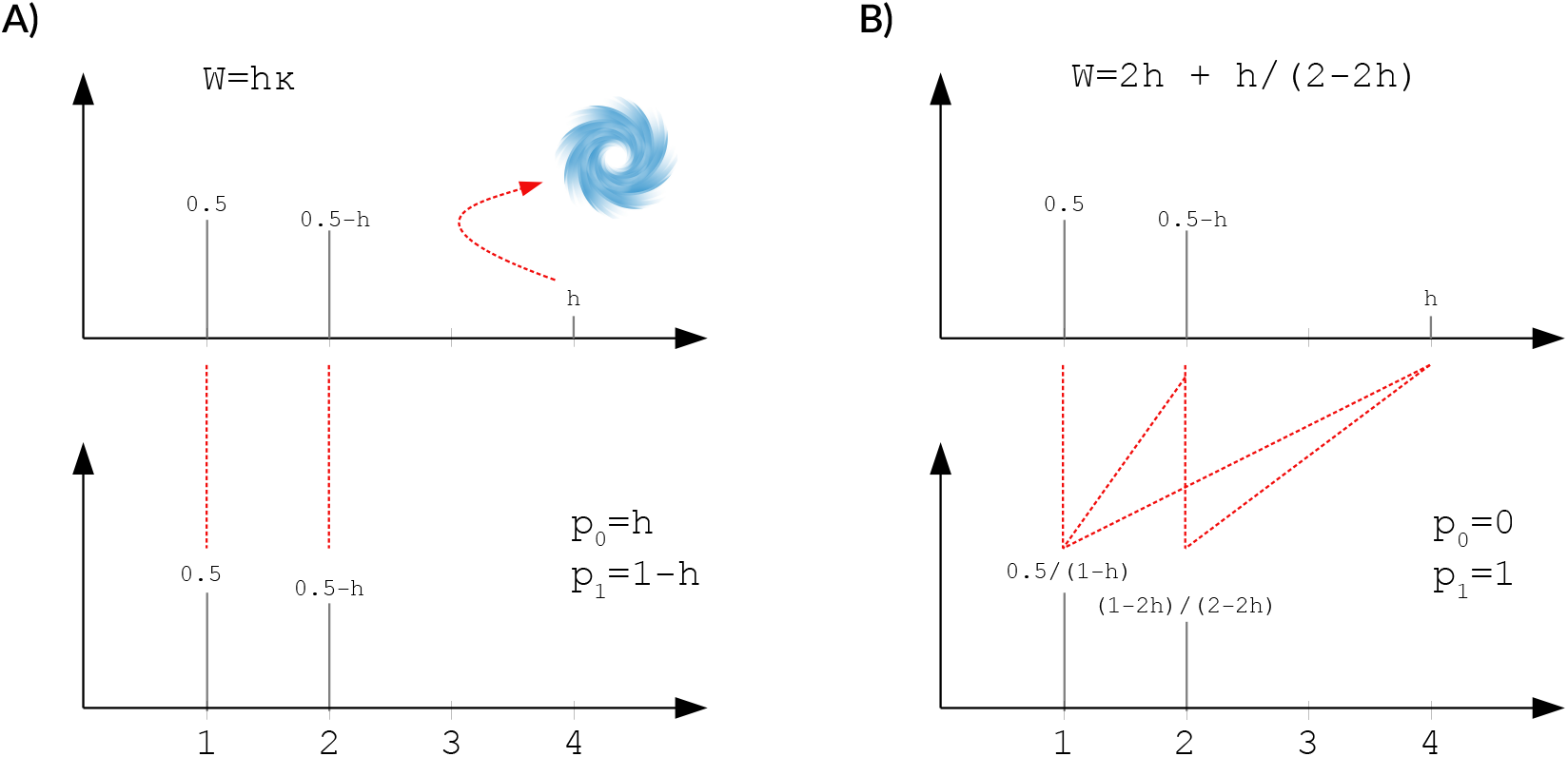
An example of deconvolution when *κ* controls the maximum transport distance. In this example, setting *κ* < 2 makes the scenario A less costly than scenario B, and prohibits the transport of the small experimental peak at 4 Da. The transport of this peak over a distance of 2 Da is permitted when *κ* > 2.

**Figure S3:**
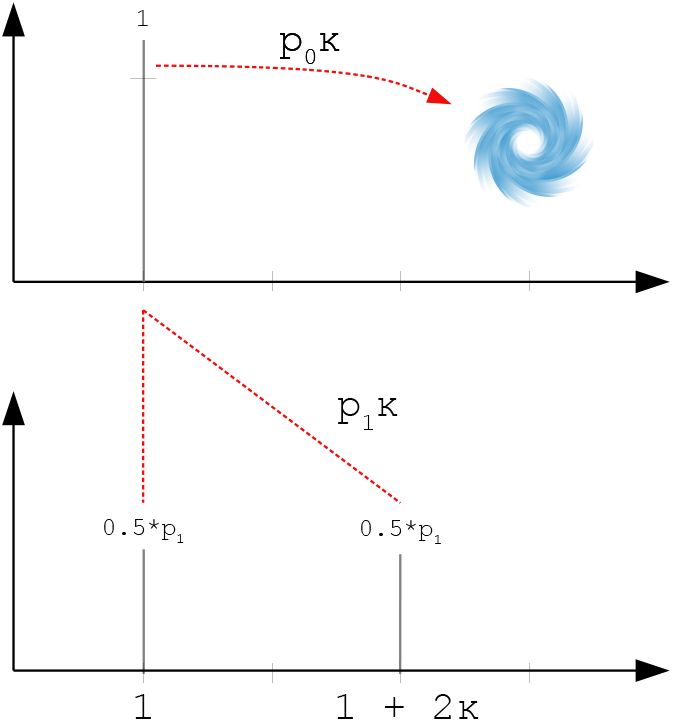
An example of a non-unique solution to the deconvolution problem. The observed spectrum is shown at the top, the theoretical one at the bottom. The Wasserstein distance is equal to *W*(*μ, v*) = *p*_0_*κ* + *p*_1_*κ* = *κ* and does not depend on the proportion of the expected spectrum *v*.

In order to solve the problem (15) under the above assumptions, we will convert it to a linear program. A linear program is a problem of finding a minimum of a linear function under a set of linear constraints. We follow the ideas of converting a Least Absolute Deviations regression problem into a linear program outlined in [19].

Recall that we write *S* = (*s*_1_, *s*_2_, … , *s*_*n*_) for a sorted list of all observed m/z values. Since the CDFs of the spectra are step functions, the integral in the problem (15) is equal to a simple sum:

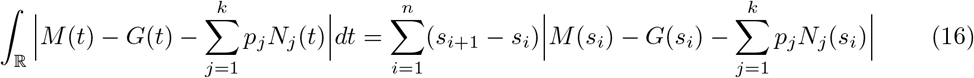

As discussed in the main text, equations of the above form admit a natural interpretation in terms of optimal transport. The difference of CDFs at the point *s*_*i*_ is the difference in the amount of ion current between the compared spectra on the left hand side of this point. This difference needs to be balanced by transporting the ion current either to or from the next point, *s*_*i*+1_. Therefore, the summands can be interpreted as the amount of ion current that flows between points *s*_*i*_ and *s*_*i*+1_ multiplied by the interval length.

Let *M*_*i*_ = *M*(*s*_*i*_), *N*_*ij*_ = *N*_*j*_(*s*_*i*_) and *G*_*i*_ = *G*(*s*_*i*_). Let *l*_*i*_ = *s*_*i*+1_ − *s*_*i*_ be the *i*-th interval length, and let 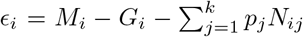 be the ion current flow between *s*_*i*_ and *s*_*i*+1_. The task now is to minimize *ϵ*_*i*_ over proportions *p*_*j*_ and amounts of removed signal *g*_*i*_. Note that, since the spectra are normalized, the ion current balances out at *s*_*n*_, so that *ϵ*_*n*_ = 0. Therefore, in the optimization problems below, we consider *ϵ*_*i*_ only for *i* = 1, 2, … , *n* − 1.

Using the above notation notation, we reformulate our optimization problem as follows:

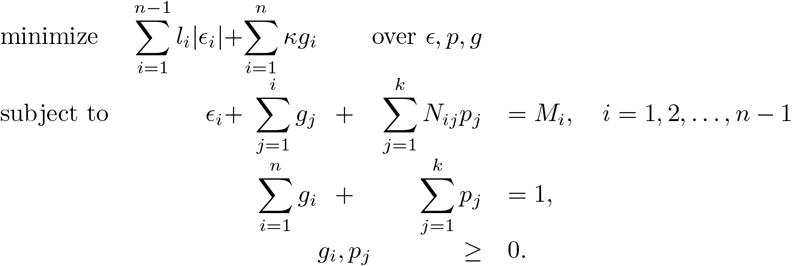

Note that while we enforce a constraint that the total amount of removed signal does not exceed the leftover signal in the observed spectrum, we do not enforce such a constraint peak-wise. That is, during the course of numerical optimization, it may happen that the proportion of signal removed from the *i*-th experimental peak will temporarily exceed the intensity of that peak. However, such a peak-wise constraint is fulfilled automatically in the optimized denoising scheme.

The objective function, 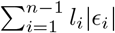, is not yet linear. However, by splitting the error *ϵ*_*i*_ into a positive part, 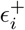, and a negative part, 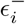, so that 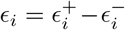, we can rewrite the above optimization problem as a linear program:

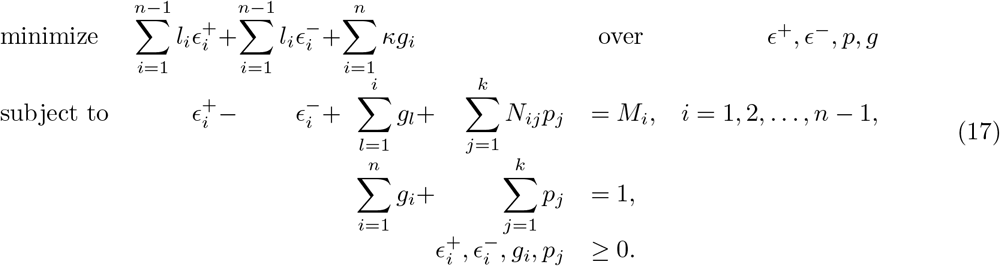

The above linear program has 3*n* − 2 + *k* variables and *n* + 1 constraints. For large spectra, where *n* can be of the order of tens of thousands peaks, this leads to a computationally intensitve optimization problem. In order to obtain a more efficient algorithm, we consider a *dual* problem. A comprehensive treatment of the duality theory in linear programming can be found in [18]. In short, each linear program admits a so-called dual program which is equivalent in the sense that it has the same optimal value of the optimized function. Furthermore, after solving the dual program, we can easily reconstruct the optimal values of the variables of the *primal* program (17).

The dual formulation of the program (17) is as follows:

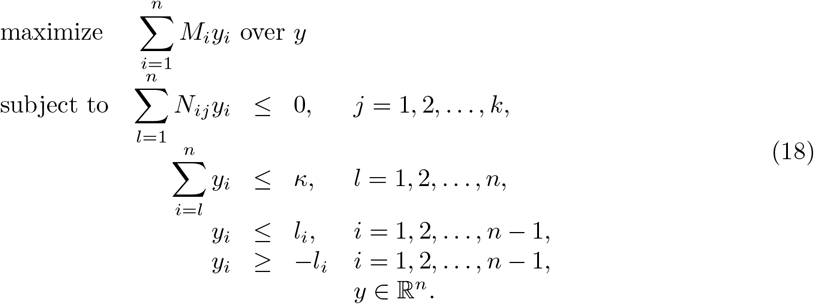

In the above dual program, we have *n* variables, *n* + *k* constraints and 2*n* − 2 bounds. We now proceed to further simplify the program.

Let *U* = ([*i* ≥ *j*])_*i,j*=1,2,…,*n*_ be an *n* × *n* square, lower-triangular binary matrix with ones on and below the diagonal and zeros above it. We use it to re-write the above dual program in matrix notation:

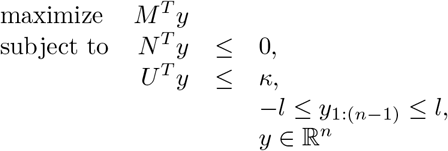

Let *W* = (*v*_*j*_(*s*_*i*_)) be the matrix of intensities of the theoretical spectra on the points *s*_*i*_ for *i* = 1, 2, … , *n*, i.e. the theoretical spectra stacked column-wise. Similarly, let *V* = (*μ*(*s*_*i*_)) be a vector of intensities of experimental spectrum for *i* = 1, 2, … , *n*. Note that we have *N* = *UW* and *M* = *UV* , and substitute that into the program formulation:

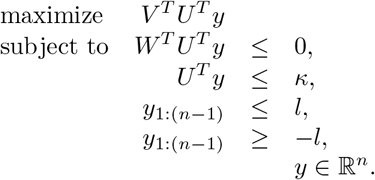

Since the matrix *U* is full-rank, substitution *z* = *U*^*T*^_*y*_ is a valid variable change. Note that

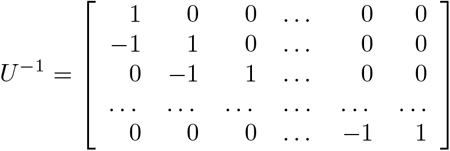

and since *y* = *U*^−T^_*z*_, we have *y*_*i*_ = *z*_*i*_ − *z*_*i*+1_ for *i* = 1, 2, … , *n* − 1 and *y*_*n*_ = *z*_*n*_. After substituting and rearranging rows of the linear program we obtain

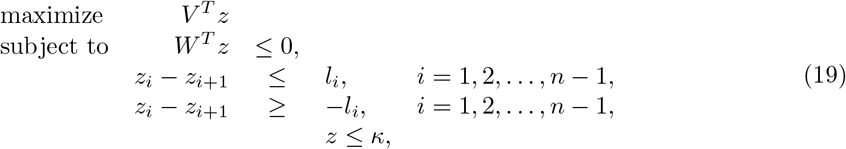

which is a program with *n* upper-bounded variables and *k* + 2*n* − 2 constraints.

Program (19) is the dual linear program in its final form. Its major advantage over the first representation of the dual program is the sparsity of matrix *W* as opposed to matrix *N* , meaning that many of its values are zero. Modern implementations of the Simplex algorithm, one of the algorithms used to solve linear programs, take advantage of matrix sparsity to speed up the computations.

Since the linear program (19) was obtained by rearranging the lefthand side terms in Program (18), the optimal target function value stays unchanged, just as the values of the primal program variables corresponding to the constraints. The latter can be obtained from the solution, as the modern Simplex implementations implicitly solve both the dual and the primal problem. The probability values, *p*_*j*_, can be obtained as the duals to the constraints *W*^*T*^_*z*_ ≤ 0, while the noise amounts *g*_*i*_ can be obtained as the duals to the constraints *z*_*i*_ ≤ *κ*.

On a final note, note that we have allowed for non-zero noise value *g*_*i*_ on non-experimental peaks.

In the dual linear program (19), this is visible as upper bounds for all *z*_*i*_ variables, instead of just the ones that correspond to points *s*_*i*_ with a non-zero experimental signal. This does not influence the results, as non-zero *g*_*i*_ at a point without any experimental signal is always sub-optimal. If one would like to explicitly forbid non-zero values of *g*_*i*_ at masses without experimental signal, it would suffice to remove upper bounds for *z*_*i*_ corresponding to purely theoretical peaks. Removing bounds leads to a simplification of the feasible region, which speeds up the Simplex algorithm. However, we have found out that the speedup obtained this way is negligible in practice.

## 4.3 Simulation of mass spectra

To simulate random molecular formulas, we use Algortihm 1 for *ε* = (*C, O, N, S, P*) and = (12, 16, 14, 32, 31). For each element, the number of atoms is sampled uniformly from 0 to the maximum number allowed by the remaining mass, and the remaining mass is filled with hydrogen atoms. Note that, in this algorithm, the order of elements influences their abundance. Elements which are closer to the beginning of the list *ε* tend to be more abundant that those at the end of the list.

**Algorithm 1:**
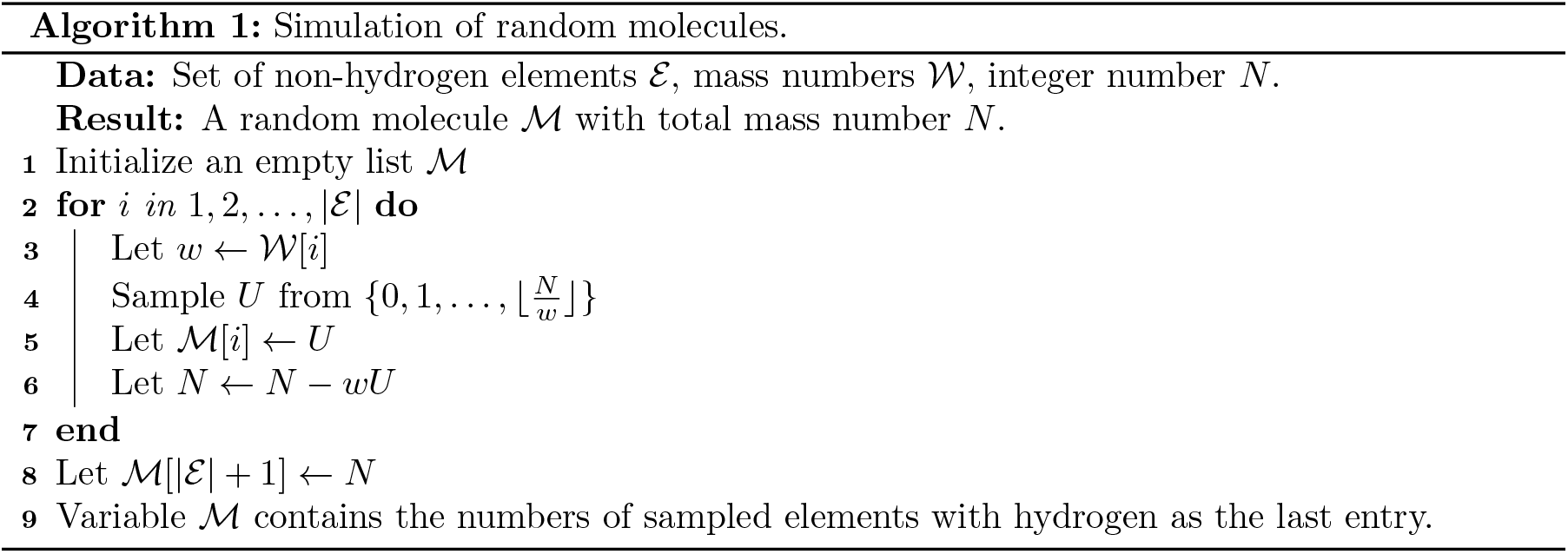
Simulation of random molecules.

We have simulated 100 sets of molecular formulas (referred to as replicates) for each combination of the following parameters:

- Nominal mass of molecules: 60, 120, 600, 1200, 6000, 12000,
- Number of overlapping isotopic envelopes: 1, 2, 3, 4, 5, 6, 7, 8.

After the molecules were simulated, theoretical spectra were computed using the IsoSpecPy package [20]. To construct observed spectra, we have simulated several sources of measurement distortions. An example result of the simulation is shown in Fig. S4.

First, we have simulated the effect of a finite number of molecules, which causes the peak heights to be variable due to random numbers of isotopologues. We assume that each observed spectrum is formed by *N* = 10000 ions to obtain a moderate to high variability of peak heights. For each set of molecular formulas, we sample their proportions *p*_1_, … , *p*_*k*_ uniformly from a unit simplex 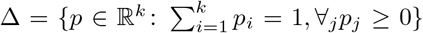. The number of ions of the *j*-th molecular formula is then equal to 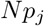. Each ion is then assigned to an isotopic composition according to the probabilities computed by IsoSpec, under the assumption of standard isotopic compositions of elements. Next, we assume that each ion contributes a random amount of signal intensity to its spectrum, with a Gaussian distribution with mean 1 and standard deviation 0.001. A mass spectrum is then obtained by summing the signal contributions of all ions.

Next, to simulate chemical noise, we have added 50 random peaks, with uniformly distributed m/z values and Gamma-distributed intensities (shape=scale=2). Those peaks were then scaled so that the total amount of noise had a Beta distribution (alpha=1.444, beta=5). The parameters were selected so that, on average, the noise peaks amount for 10% of the total signal intensity in the spectrum. Further distortions were different for centroided and profile observed spectrum.

In the case of simulated centroided spectra, we additionally simulate the effects of limited resolution and accuracy and errors introduced during the preprocessing procedures such as peak picking. To each m/z value we add a Gaussian random variable with mean 0 and standard deviation of 0.002. The parameters were chosen to obtain only two accurate decimal digits in the m/z values. To simulate a limited resolving power, we round the m/z to three decimal digits and merge peaks with equal masses.

In the case of simulated profile spectra, we simulate the effect of limited resolving power and electronic noise. We use a Gaussian filter with a standard deviation of 0.0025 to obtain a resolving power of 100 000 at 600 Da. Next, to each intensity measurement we add a Gaussian random variable with mean zero and standard deviation of 0.0001.

## 4.4 Interpolation and centroiding

In this subsection, we show pseudo codes for additional algorithms used in this work to perform linear interpolation of profile spectra before averaging, and centroid the spectra.

## Piecewise-linear interpolation of spectra

In order to obtain an average profile spectrum, we need to ensure that all spectra share a common set of m/z values. However, those values often slightly differ between spectra. In order to correct that, we interpolate each spectrum by a piecewise linear function, so that the interpolated intensity in a given point is a weighted average of the neighboring measured intensities.

Although the terminology may sound obscure, the idea behind the piecewise-linear interpolation is straightforward: we join pairs of consecutive points by straight lines and use those lines to approximate the signal intensities at any given set of points at the mass axis. Algorithm 2 shows a numerically optimized way to achieve this. The idea behind the algorithm is visualized in Fig. S5.

**Algorithm 2:**
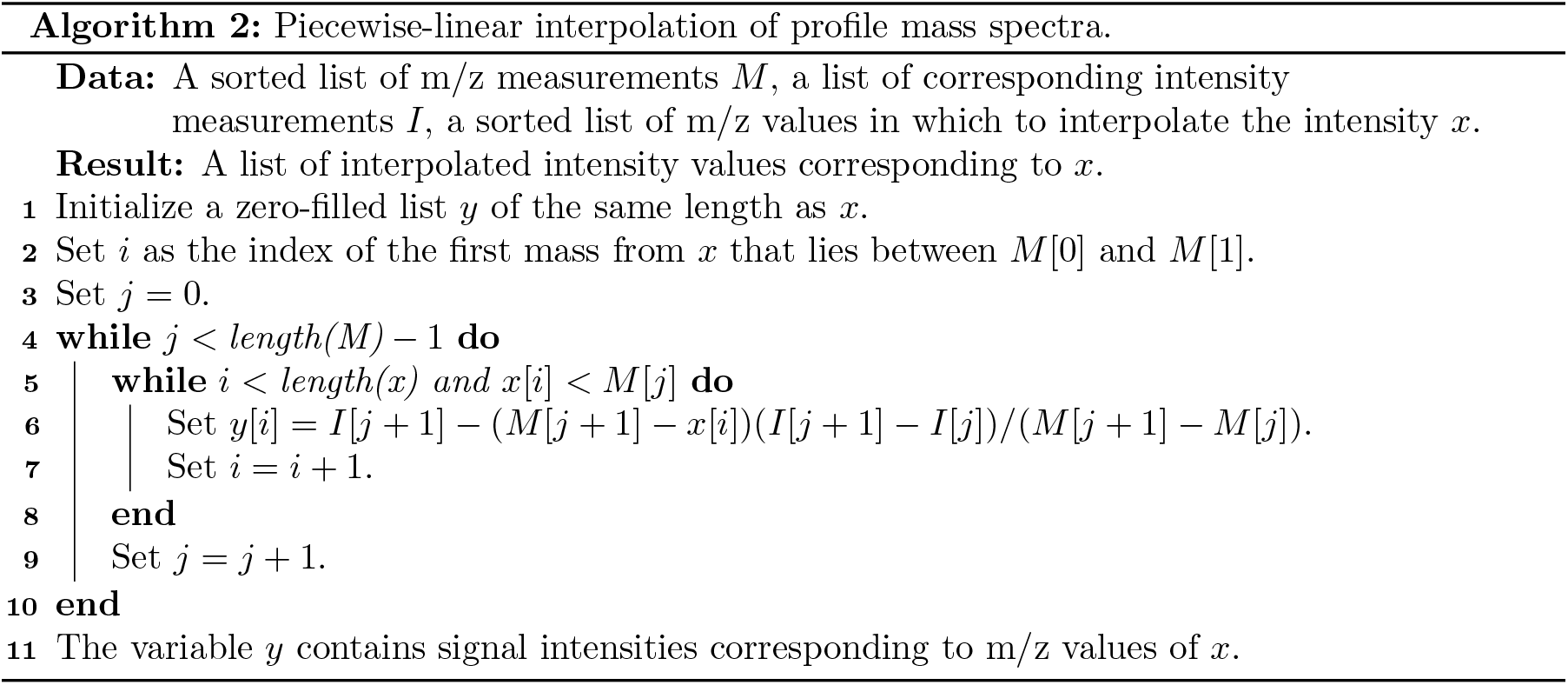
Piecewise-linear interpolation of profile mass spectra.

In our implementation, we add another constraint that if the distance between the m/z value at which we interpolate and any of it’s neighboring points is further than a given threshold, the interpolated intensity is set to 0. This is done in order to handle broad regions with no recorded intensity present in many experimental spectra.

We have used linear interpolation mainly for two tasks. The first one was to compute an average profile spectrum from the dataset of 200 spectra used in our experiments. We have set a mass axis and approximated the signal intensity of each profile spectrum at each m/z value of the axis. Next, for each m/z value the approximated intensities were summed over the 200 spectra. The second task was to obtain a constant spacing between consecutive signal measurements in profile spectra. To this end, we have set a mass axis with constant spacing and interpolated each profile spectrum at the points of the axis. The resulting spectra were subsequently subjected to deconvolution without any further processing.

## Centroiding the profile spectra

There are numerous available algorithms for peak centroiding, both open-source and proprietary. We have decided not to use the latter in this work, as it is not possible to determine the precise way they work. Instead, we have implemented a simple algorithm that detects local maxima of intensity and integrates peaks within regions delineated by an intensity threshold, expressed as a proportion of the apex intensity.

**Figure S4:**
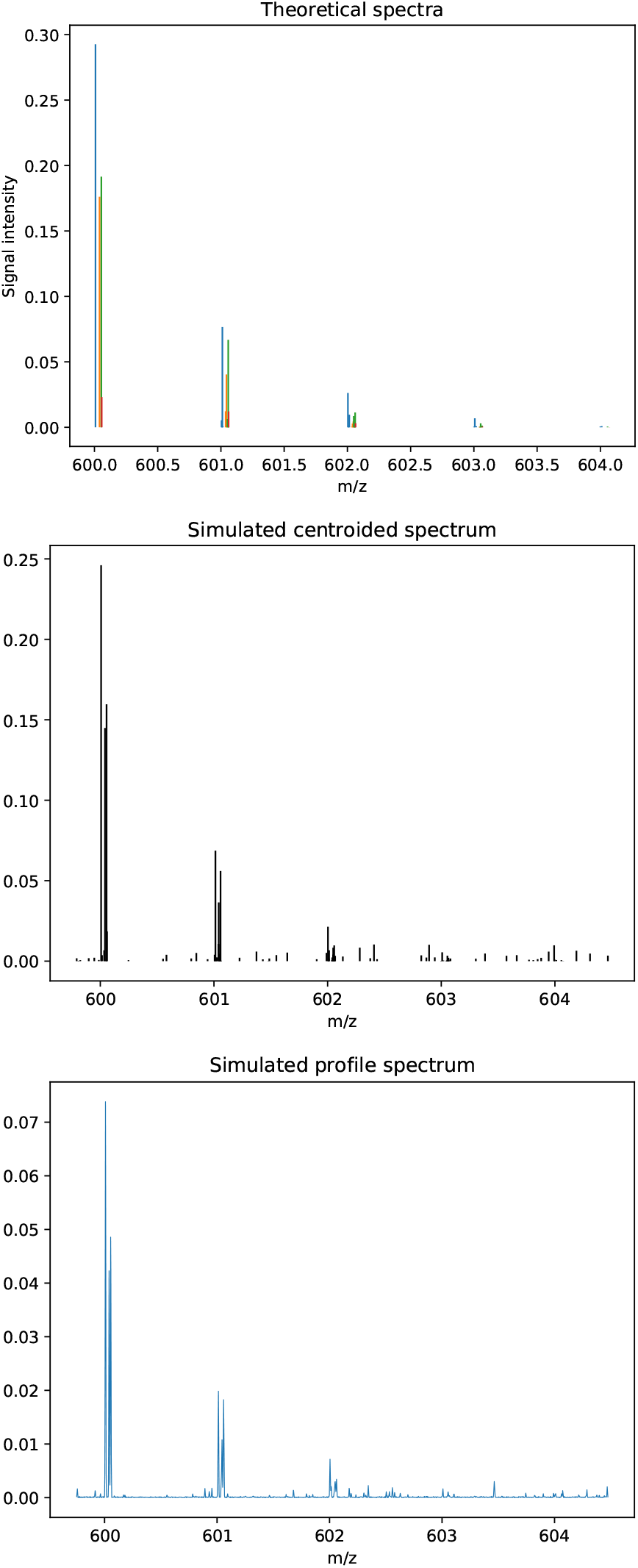
An example of simulated mass spectra of four overlapping isotopic envelopes. The spectrum on the left was used to construct a centroided and a profile observed spectrum with measurement distortions. Isotopic envelopes of different ions are highlighted with different colors.

**Figure S5:**
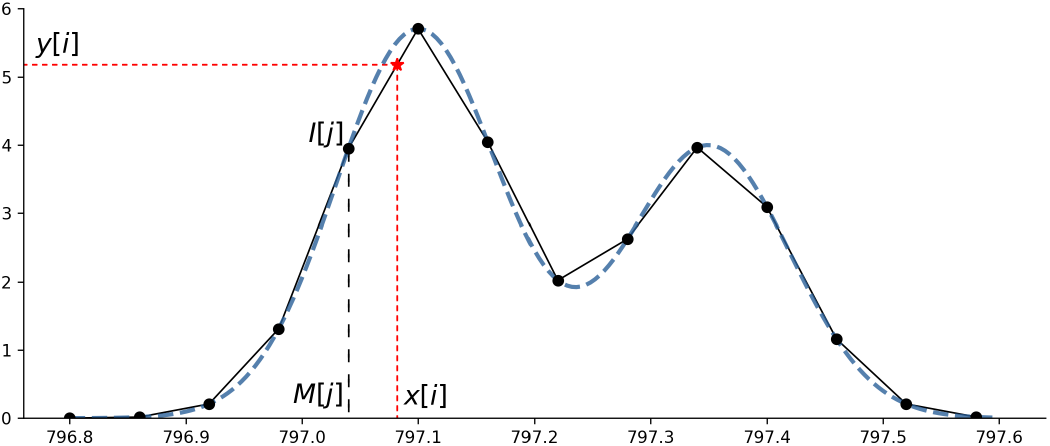
A graphical description of piecewise-linear interpolation of profile mass spectra. The blue dashed line shows the true, unobserved signal intensity. The black points show the intensity measurements *I*[*j*], corresponding to masses *M*[*j*], observed in a profile mass spectrum. The black lines show the interpolated intensities. The red dashed lines show a result of approximation of the true signal, *y*[*i*], at the point *x*[*i*]. The indexes *i* and *j* correspond to the ones used in Algorithm 2. Note that in actual spectra the measured points are more densely spaced, resulting in a much better interpolation.

Note that, after peak location is determined, there are two main approaches to obtain peak intensity: either as the apex intensity, or as the peak area. In the context of this manuscript, only the latter is applicable, as the peak width increases with the m/z value.

**Algorithm 3:**
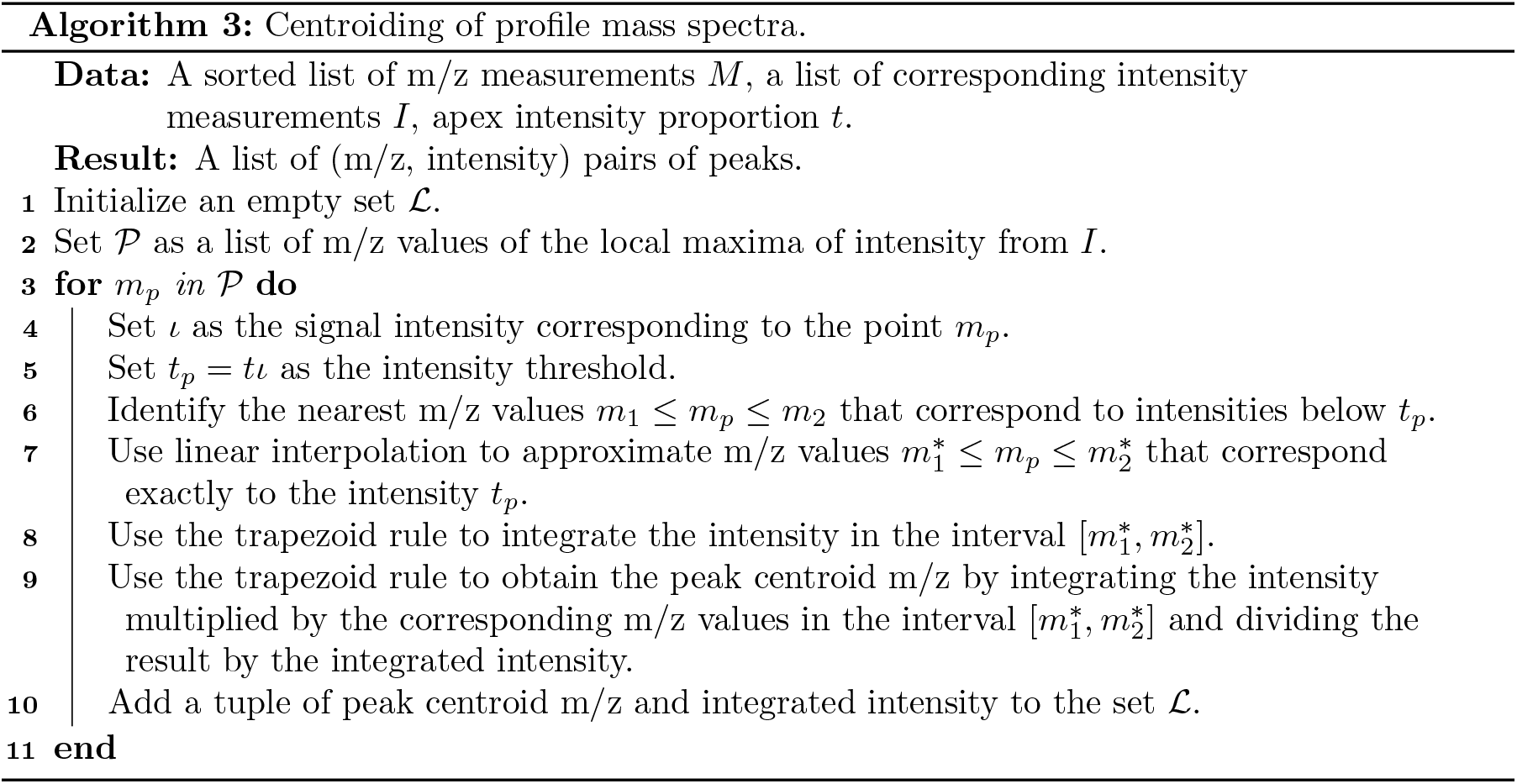
Centroiding of profile mass spectra.

In our implementation, we set additional constraint on maximal peak width equal to 0.5 Da. If the width of the region in which the intensity is to be integrated exceeds this threshold, the peak is discarded.

Note that, when two peaks overlap, they may share their integration region. In that case, the centroided m/z and intensities of such peaks are identical. Since we keep a set of peaks instead of a list (note the line 1 of Algorithm 3), such a peak cluster is represented by one peak. This essentially merges highly overlapping peaks into one. Separating highly overlapping peak clusters requires much more sophisticated approaches (see e.g. [23]).

**Table S1:**
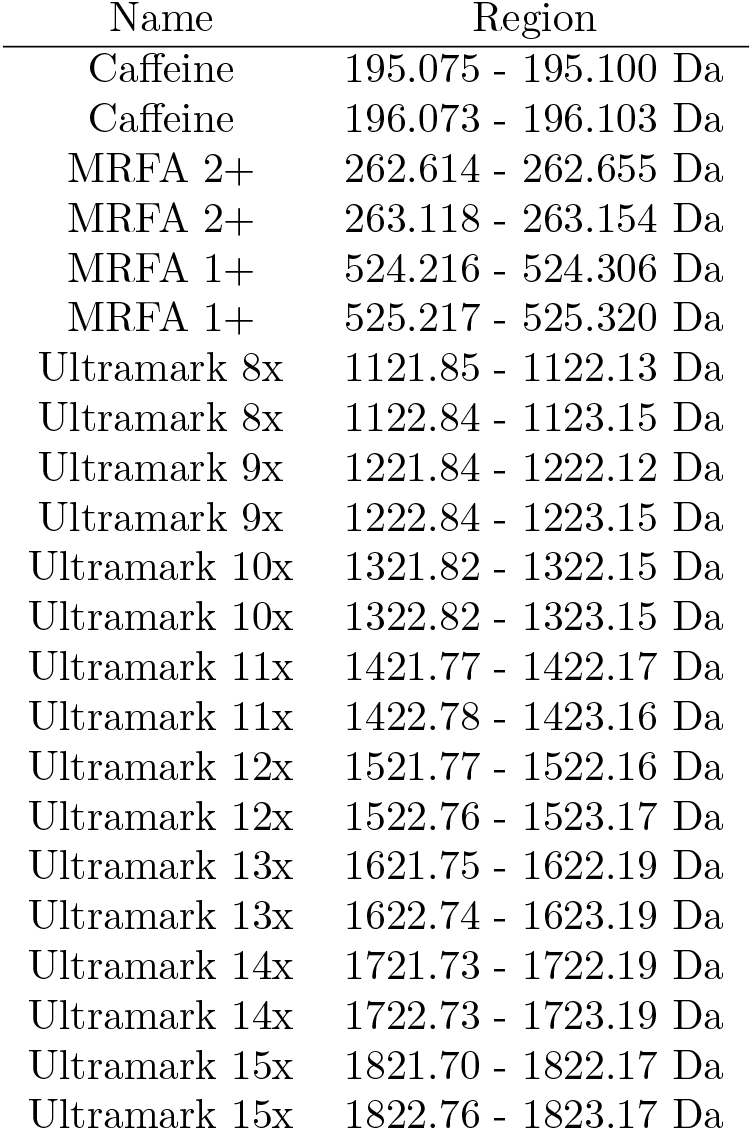
Regions of the m/z axis used to compute ion signal intensities. Ultramark 8x denotes Ultramark 1621 with 8 CF_2_CF_2_ groups, etc.

In our implementation, when we identify the integration region in line 6 of Algorithm 3, we additionally require that the intensity is monotonically decreasing with the distance from the apex. When we detect that the signal intensity starts to increase, we discard the peak. This ensures that only the highest peak of any peak cluster is considered. It also allows to discard numerous small peaks that occur due to background noise.

## Supplementary figures and tables

**Figure S6:**
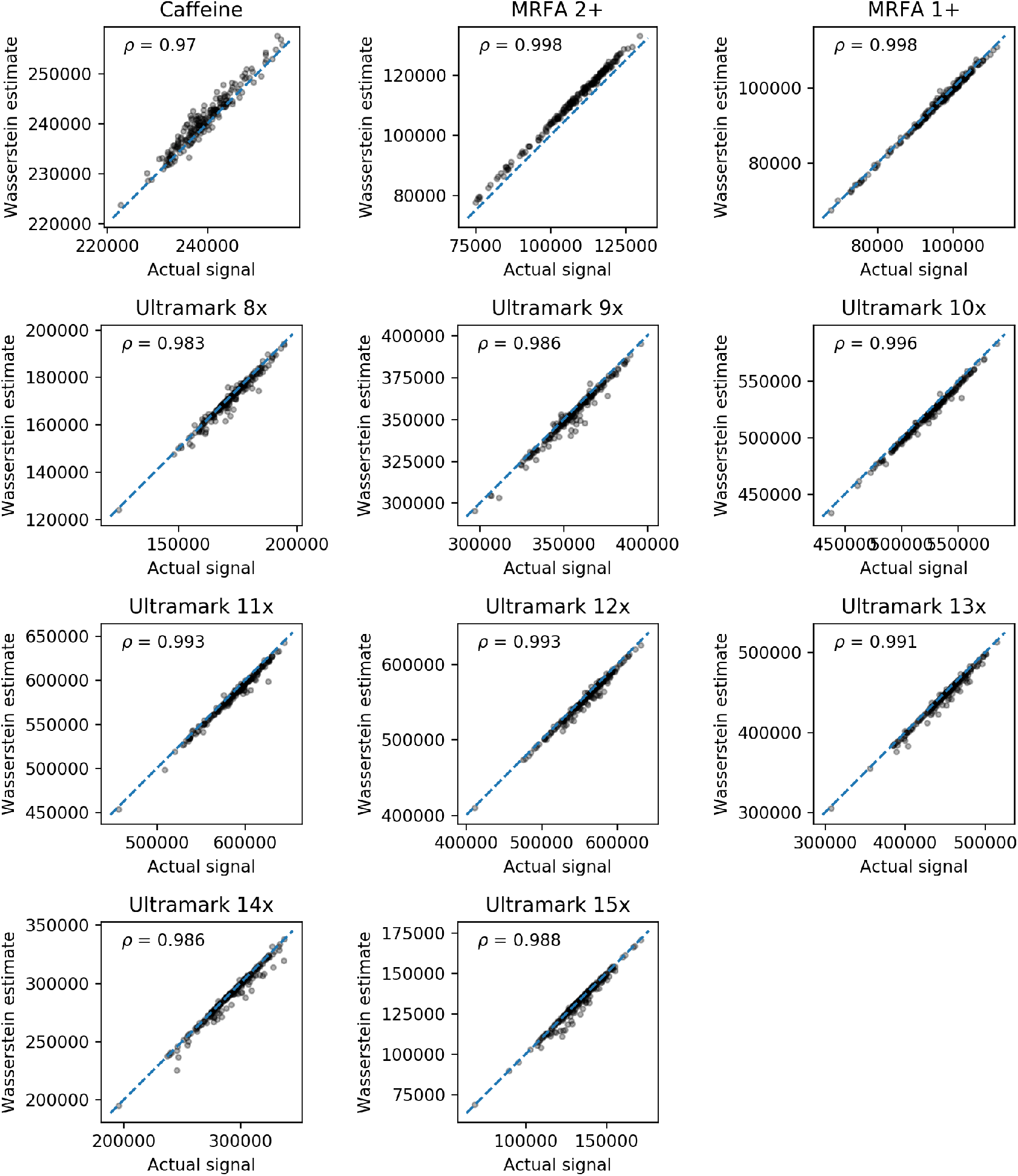
The deconvolution results for 200 centroided spectra of Pierce LTQ Velos ESI Positive Ion Calibration Solution (Thermo Scientific). The plots show ion intensities estimated by the linear deconvolution based on the Wasserstein distance versus manually integrated peak areas. Each point corresponds to a mass spectrum. Numbers in top-left corners represent the Pearson correlation. Note the different scales in plots, due to different average signal intensities of different molecules.

**Figure S7:**
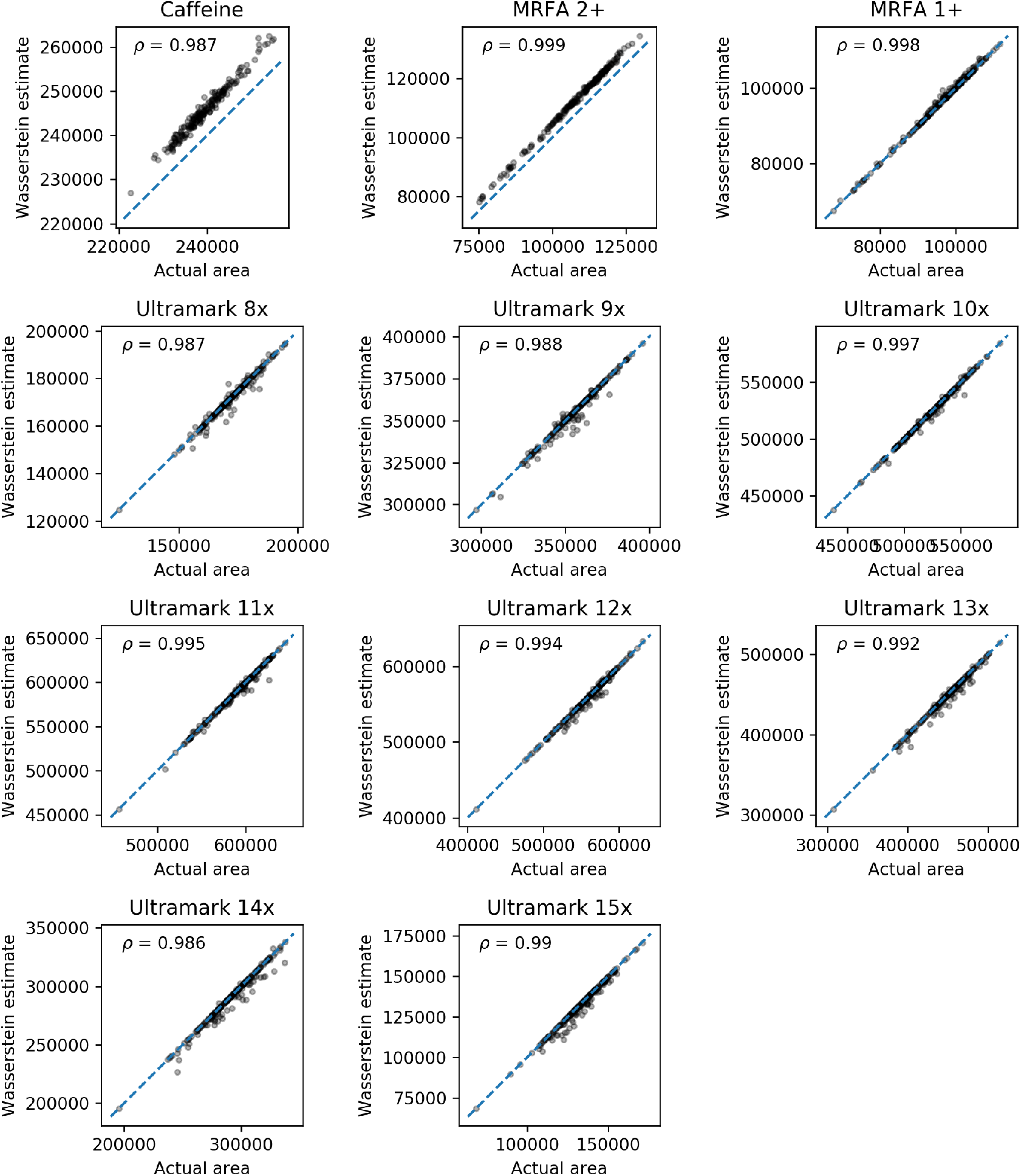
The deconvolution results for 200 profile spectra of Pierce LTQ Velos ESI Positive Ion Calibration Solution (Thermo Scientific). The plots show ion intensities estimated by the linear deconvolution based on the Wasserstein distance versus manually integrated peak areas. Each point corresponds to a mass spectrum. Numbers in top-left corners represent the Pearson correlation. Note the different scales in plots, due to different average signal intensities of different molecules.

**Figure S8:**
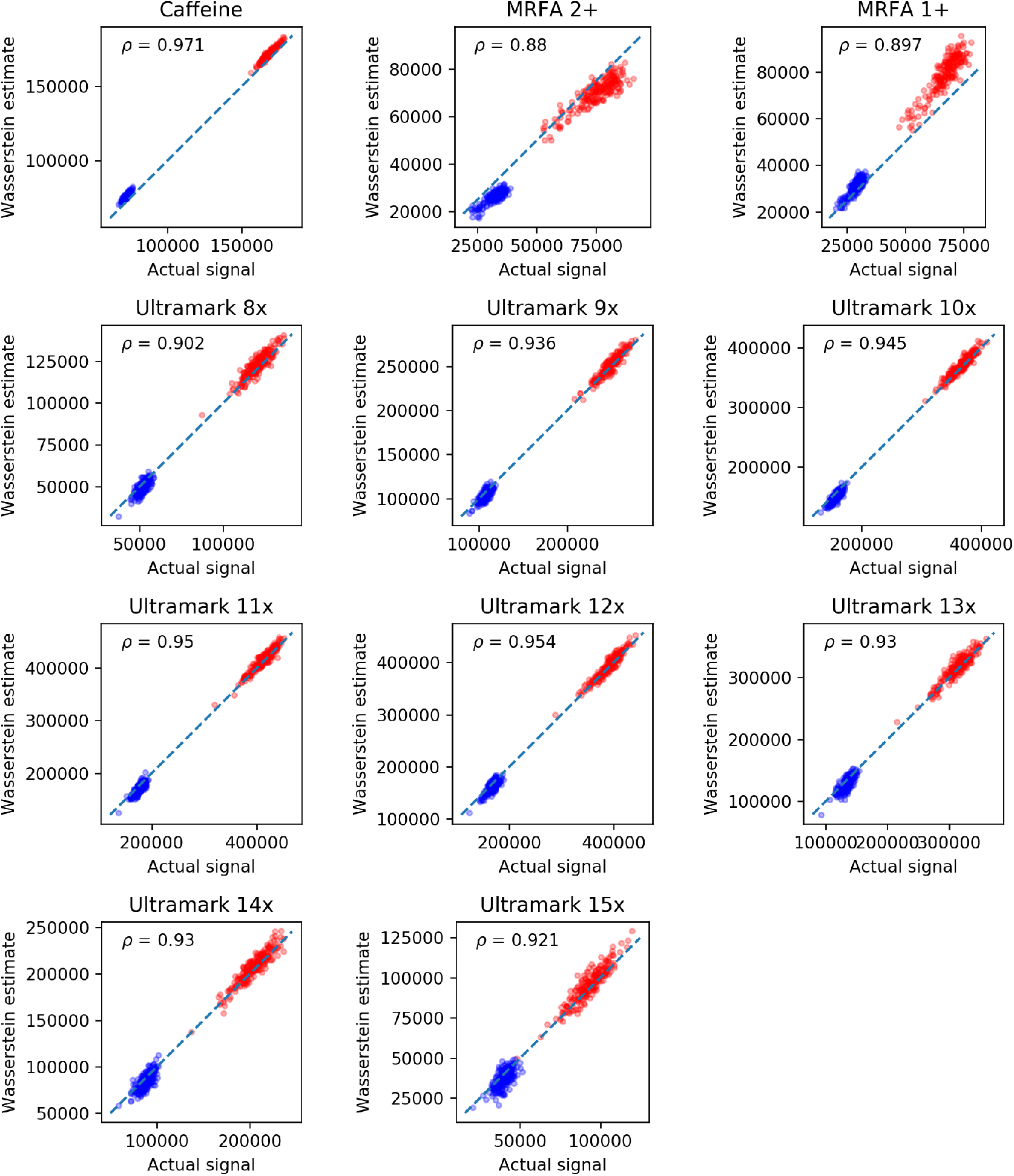
The deconvolution results on 200 spectra of Pierce LTQ Velos ESI Positive Ion Calibration Solution (Thermo Scientific) after introducing overlapping isotopic envelopes. Red: signals from original spectra; Blue: signals from spectra shifted by one hydrogen mass. Note the different scales on the X and Y axes.

**Figure S9:**
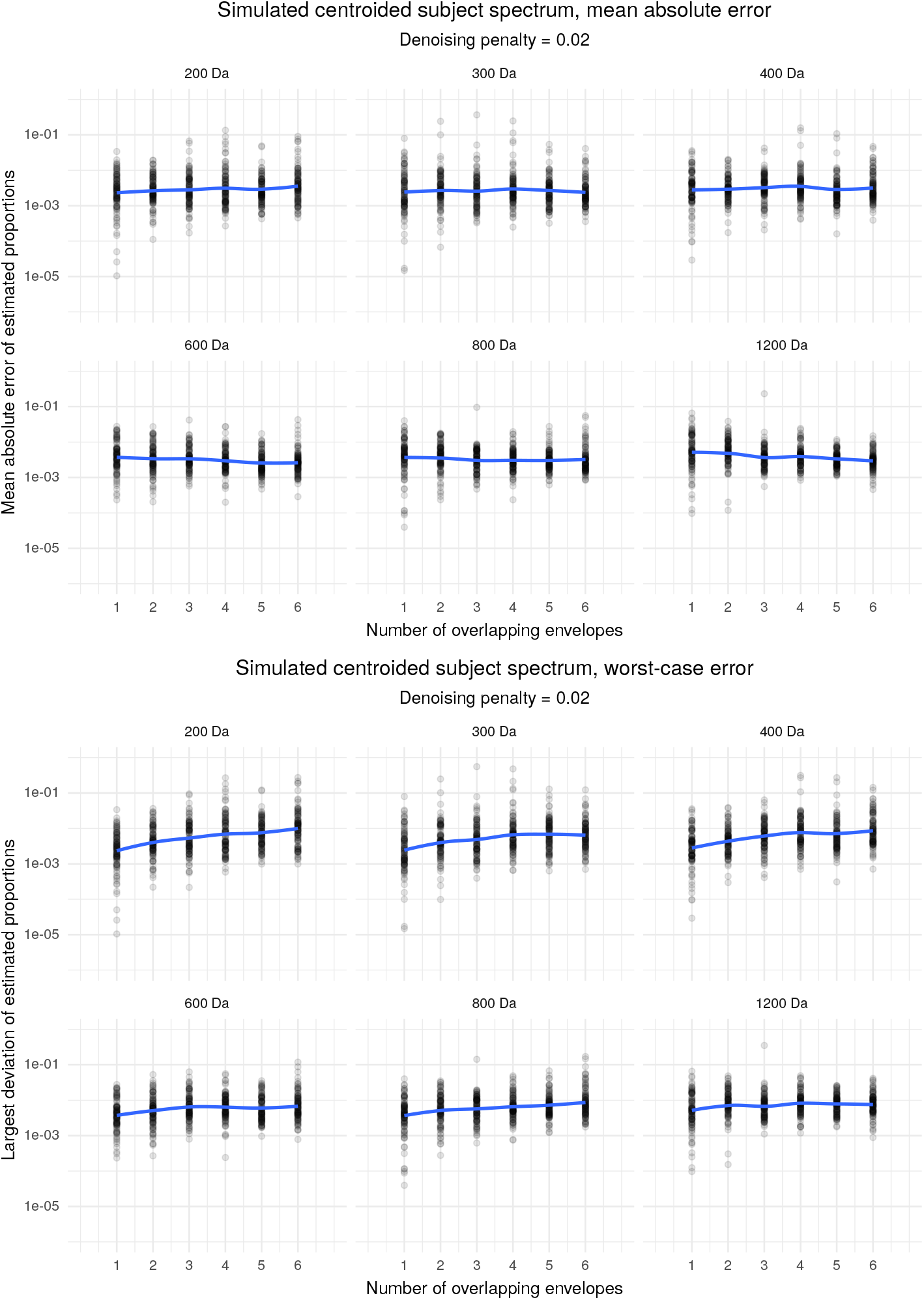
Errors of estimation of molecule proportions on centroided experimental (i.e. subject) spectra. Top: mean absolute deviation of estimation per spectrum. Bottom: largest deviation of estimation per spectrum. Note the logarithmic scale of the plots.

**Figure S10:**
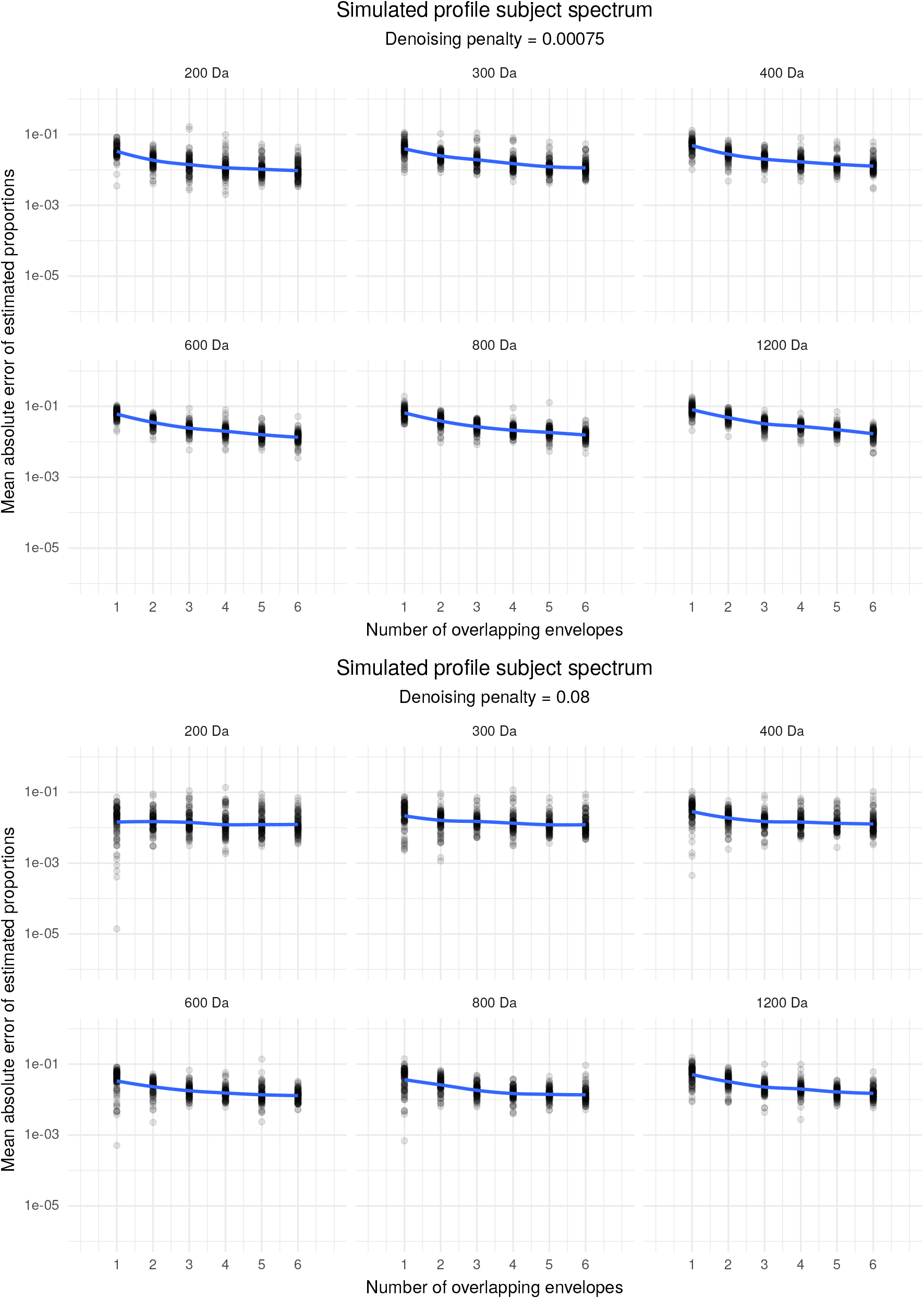
Mean absolute deviation of estimation of molecule proportions on profile experimental (i.e. subject) spectra for two denoising penalties. Note the logarithmic scale of the plots.

